# Real-time dynamics of *Plasmodium* NDC80 reveals unusual modes of chromosome segregation during parasite proliferation

**DOI:** 10.1101/767830

**Authors:** Mohammad Zeeshan, Rajan Pandey, David J.P. Ferguson, Eelco C. Tromer, Robert Markus, Steven Abel, Declan Brady, Emilie Daniel, Rebecca Limenitakis, Andrew R. Bottrill, Karine G. Le Roch, Anthony A. Holder, Ross F. Waller, David S. Guttery, Rita Tewari

**Affiliations:** School of Life Sciences, Queens Medical Centre, University of Nottingham, Nottingham, NG7 2UH, UK; Nuffield Department of Clinical Laboratory Science, University of Oxford, John Radcliffe Hospital, Oxford, OX3 9DU, UK; Department of Biological and Medical Sciences, Faculty of Health and Life Science, Oxford Brookes University, Gipsy Lane, Oxford OX3 0BP, UK; Department of Biochemistry, University of Cambridge, Cambridge, CB2 1QW, UK; Department of Molecular, Cell and Systems Biology, University of California Riverside, Riverside, California, United States of America; Institute of Cell Biology, University of Bern, Bern 3012, Switzerland; School of Life Sciences, Gibbelt Hill Campus, University of Warwick, Coventry, CV4 7AL, UK; Malaria Parasitology Laboratory, The Francis Crick Institute, London, NW1 1AT, UK; Leicester Cancer Research Centre, University of Leicester, Leicester, LE2 7LX, UK

**Keywords:** Malaria, *Plasmodium*, kinetochore, NDC80 complex, chromosome segregation, atypical cell division, endomitosis, meiosis

## Abstract

Eukaryotic cell proliferation requires chromosome replication and precise segregation to ensure daughter cells have identical genomic copies. The genus *Plasmodium*, the causative agent of malaria, displays remarkable aspects of nuclear division throughout its lifecycle to meet some peculiar and unique challenges of DNA replication and chromosome segregation. The parasite undergoes atypical endomitosis and endoreduplication with an intact nuclear membrane and intranuclear mitotic spindle. To understand these diverse modes of *Plasmodium* cell division, we have studied the behaviour and composition of the outer kinetochore NDC80 complex, a key part of the mitotic apparatus that attaches the centromere of chromosomes to microtubules of the mitotic spindle. Using NDC80-GFP live-cell imaging in *Plasmodium berghei* we observe dynamic spatiotemporal changes during proliferation, including highly unusual kinetochore arrangements during sexual stages. We identify a very divergent candidate for the SPC24 subunit of the NDC80 complex, previously thought to be missing in *Plasmodium*, which completes a canonical, albeit unusual, NDC80 complex structure. Altogether, our studies reveal the kinetochore as an ideal tool to investigate the non-canonical modes of chromosome segregation and cell division in *Plasmodium.*

**Summary Statement:** The dynamic localization of kinetochore marker NDC80 protein complex during proliferative stages of the malaria parasite life cycle reveals unique modes of chromosome segregation.

## Introduction

Mitosis and meiosis are fundamental processes in cell division that enable DNA replication and chromosome segregation, and allow eukaryotic organisms to proliferate, propagate and survive. During these processes, microtubular spindles form to facilitate an equal segregation of duplicated chromosomes to the spindle poles. Chromosome attachment to spindle microtubules (MTs) is mediated by kinetochores, which are large multi-protein complexes assembled on centromeres located at the constriction point of sister chromatids (Cheeseman, 2014; McKinley and Cheeseman, 2016; Musacchio and Desai, 2017; Vader and Musacchio, 2017). Each sister chromatid has its own kinetochore, oriented to facilitate movement to opposite poles of the spindle apparatus. During anaphase, the spindle elongates and the sister chromatids separate, resulting in segregation of the two genomes during telophase. The NDC80 complex is the major component of the kinetochore and mediates its attachment to spindle MTs. In most model organisms, it is a member of the network of conserved Knl1, Mis12 and NDC80 complexes (KMN) (McKinley and Cheeseman, 2016; Petrovic et al., 2016). The ∼170-190 kDa NDC80 complex has two globular domains at either end of a ∼57 nm elongated coiled-coil, forming a dumb-bell shape. It is a heterotetramer comprising a 1:1:1:1 ratio of NDC80 (also known as HEC1 in humans), NUF2, SPC24 and SPC25 sub-complexed as two heterodimers: NDC80 with NUF2 and SPC24 with SPC25 (Ciferri et al., 2005; Farrell and Gubbels, 2014; Wei et al., 2005). The C-terminal end of the SPC24-SPC25 dimer anchors the complex to the kinetochore; whereas the NDC80-NUF2 dimer mediates plus-end MT binding through its calponin homology domain (CHD) (Alushin et al., 2010; Farrell and Gubbels, 2014; Sundin et al., 2011).

Malaria, caused by the apicomplexan parasite *Plasmodium* spp., remains one of the most prevalent and deadly infectious diseases worldwide, with 219 million clinical cases and 435,000 deaths in 2017 (WHO, 2018). *Plasmodium* has several morphologically distinct proliferative stages during its life cycle that alternates between vertebrate host and mosquito vector (**Fig.1**) (Francia and Striepen, 2014; Sinden, 1991a; Sinden, 1991b). A malaria parasite-infected female anopheles mosquito inoculates haploid sporozoites into the mammalian host during a blood meal. Sporozoites travel through the blood stream to the liver and infect hepatocytes, where the parasite replicates and develops into a multinucleated schizont. At the end of this exo-erythrocytic schizogony the host cell is ruptured to release haploid merozoites, which infect erythrocytes. In the intra-erythrocytic phase, an initial ring stage form develops into a trophozoite and then into a schizont where multiple rounds of asexual multiplication occur (erythrocytic schizogony). At the end of schizogony, host cell rupture releases further merozoites that infect new erythrocytes.

**Fig. 1:**
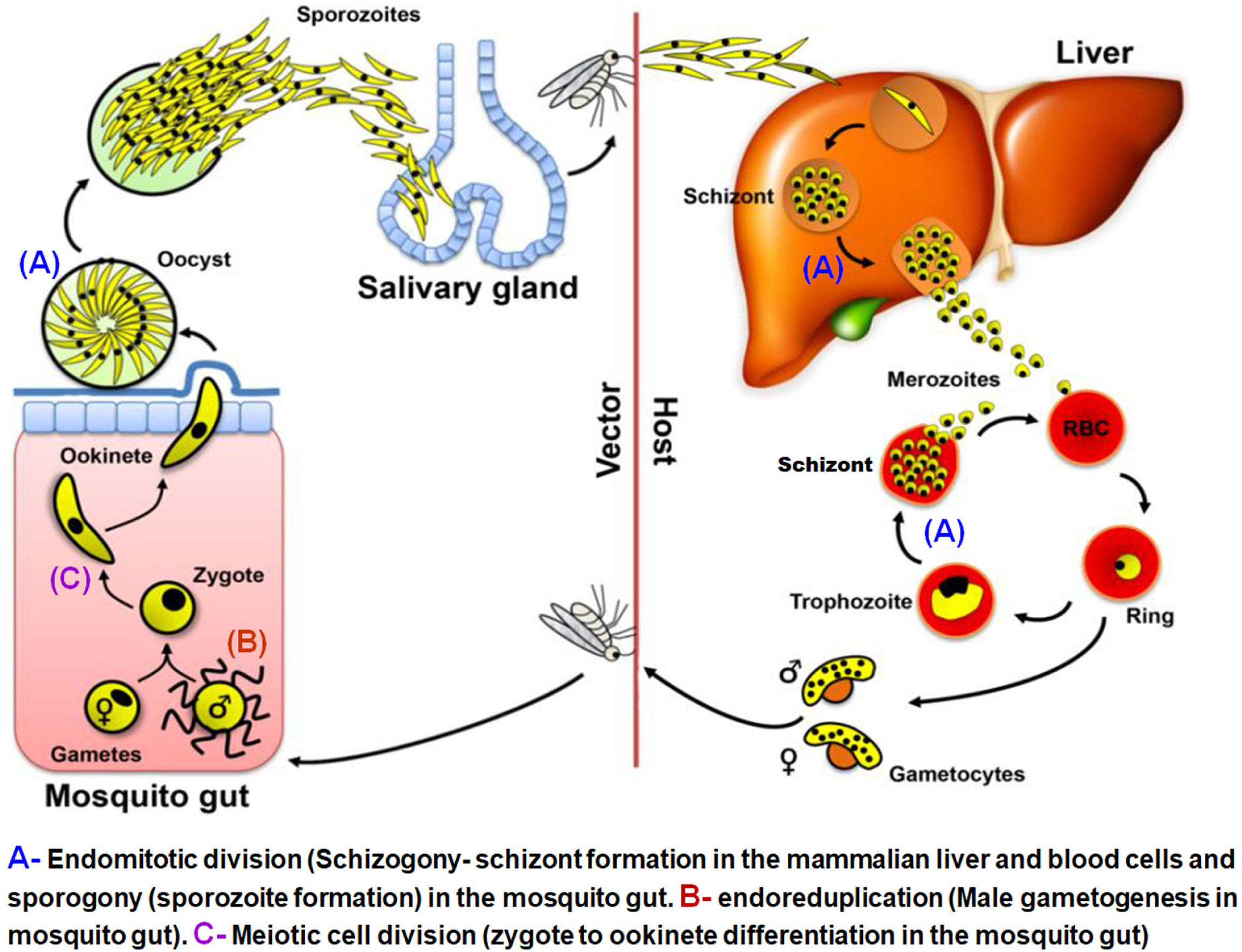
Life cycle of rodent malaria parasite *Plasmodium berghei*. (**A)** represents endomitotic division: schizogony (schizont formation) in the liver and blood cells of the mammalian host and sporogony (sporozoite formation) in the mosquito gut. (**B)** and **(C)** represent atypical mitotic division by endoreduplication during male gametogenesis and meiotic division during the zygote to ookinete differentiation in the mosquito gut, respectively.

Following erythrocyte invasion, some parasites differentiate into male (micro) and female (macro) gametocytes to initiate the sexual phase of the life cycle, which occurs inside the mosquito. These haploid parasites are arrested at the G0/G1 phase of the cell cycle (Arnot and Gull, 1998). Ingestion by a mosquito activates gametogenesis. Male gametogenesis is very rapid with three rounds of genome replication from 1N to 8N and the release of eight motile haploid microgametes within 15 min. The activated female gametocyte rounds up and the macrogamete egresses from the red blood cell. Gametogenesis can be studied *in vitro* using a culture medium that mimics the mosquito midgut environment (Billker et al., 1998; Tewari et al., 2005).

After fertilisation the zygote differentiates into a motile ookinete. The ookinete invades the mosquito midgut wall where it develops into an oocyst. At this stage multiple rounds of endomitotic division occur in a process similar to schizogony, which is followed by cytokinesis to form thousands of motile sporozoites (Francia and Striepen, 2014; Gerald et al., 2011). The sporozoites are released from the oocyst and migrate to the mosquito’s salivary glands for transmission to the vertebrate host.

The life cycle of *Plasmodium* is characterized by two unique processes of mitosis and a single stage of meiosis. The first mitotic process occurs during schizogony within mammalian hepatocytes and erythrocytes, and during sporogony in oocysts in the mosquito (Sinden, 1991a; Sinden, 1991b) (**Fig. 1 A**). This mitotic division is atypical, for example no clear G2 cell cycle phase has been observed during schizogony (Arnot and Gull, 1998; Doerig et al., 2000). Furthermore, this asexual proliferation is characterised by multiple rounds of asynchronous nuclear division without chromosome condensation and in the absence of cytokinesis. Mitosis is closed, occurring without dissolution and reformation of the nuclear envelope, and the spindle-pole body (SPB)/microtubule-organising centre (MTOC), also known as the centriolar plaque (Arnot et al., 2011; Francia et al., 2015; Sinden, 1991a), is embedded within the nuclear membrane. The asynchronous nuclear division precedes cell division, leading to a multinucleate syncytium. The last round of nuclear division in these cells is synchronous and it is only after this final round of mitosis that cytokinesis occurs to form the haploid daughter merozoites or sporozoites, respectively.

The second type of mitotic division occurs during male gametogenesis following activation in the mosquito midgut (**Fig. 1 B)**. Three rounds of rapid genome duplication (from haploid to octoploid) without concomitant nuclear division (endoreduplication) are followed by chromosome condensation and nuclear budding into the male gametes during exflagellation, all within 12 to 15 min of activation (Arnot and Gull, 1998; Janse et al., 1988; Sinden, 1983). The resultant eight flagellated microgametes each contain a haploid genome (Guttery et al., 2015; Sinden et al., 2010). Fertilization of the female gamete results in a diploid zygote, which develops in the mosquito gut and differentiates over a 24-hour period into a motile ookinete (**Fig. 1 C**). It is in this stage that meiosis occurs. The DNA is duplicated once to form a tetraploid cell and then two rounds of chromosome segregation result in four discrete haploid genomes prior to nuclear division and ookinete maturity. Reductive division to haploidy presumably occurs in the subsequent oocyst during sporozoite formation (Guttery et al., 2015; Sinden, 1991a; Sinden, 1991b). Collectively, these different stages of cell division and proliferation indicate that the parasite has evolved alternate modes of chromosome replication, condensation and segregation, as well as nuclear and cell division at different stages during its life cycle.

The process of chromosome segregation and associated kinetochore dynamics, which is the key role of the mitotic apparatus throughout the life cycle, is not well understood in *Plasmodium*. To date, analysis of *Plasmodium* mitotic/meiotic spindle assembly and chromosome segregation has been performed largely using transmission electron microscopy (TEM) (Sinden et al., 1978; Sinden et al., 1976), and biochemical analysis of microtubule markers including α-tubulin (Fennell et al., 2008), and centrin associated with the putative MTOC (Gerald et al., 2011; Roques et al., 2019). An analysis of a *Plasmodium* artificial chromosome (PAC) identified a putative centromere derived from chromosome 5 (*PbCEN5*), and highlighted the dynamics of chromosome segregation during both mitotic and meiotic stages of the parasite’s life cycle (Iwanaga et al., 2010). However, there is no real-time analysis of chromosome segregation dynamics during the various proliferative stages, especially during stages that occur inside the mosquito vector. Here, we have analysed the real-time expression and spatiotemporal dynamics of NDC80, as a kinetochore marker. We generated a stable transgenic *P. berghei* line expressing NDC80 with a C-terminal GFP-tag by modifying the endogenous gene locus. Using this tool, we examined NDC80 expression and location to follow the spatiotemporal organisation of outer kinetochores during mitosis in schizogony, sporogony and male gametogenesis, and during meiosis in ookinete development. We observed unusual kinetochore dynamics as patterns of clustered foci adjacent to the nuclear DNA during endomitotic nuclear division in asexual stages, with the number of foci corresponding to the likely ploidy of individual nuclei. However, kinetochores formed an unusual bridge-like pattern during endoreduplication stages in male gametogenesis. We show there is likely a full complement of NDC80 complex subunits, with a highly divergent candidate for the previously undetected SPC24 subunit identified by a combination of proteomics and sensitive comparative sequence analysis. This transgenic parasite line expressing GFP-tagged NDC80 is a valuable resource for studying chromosome segregation and dynamics, as well as identifying and characterising the various protein complexes involved in these cell division processes.

## Results

To study kinetochore dynamics, we generated NDC80-GFP, a transgenic *P. berghei* line expressing NDC80 with a C-terminal GFP tag, by inserting an in-frame *gfp* coding sequence at the 3’ end of the endogenous *Ndc80* locus using single homologous recombination (**Fig. S1A**). To complement our GFP-based imaging studies, we also generated a NDC80-mCherry line using the same strategy (**Fig. S1A**). Successful insertion was confirmed by PCR (**Fig. S1B**). Western blot analysis of a schizont protein extract using an anti-GFP antibody revealed the NDC80-GFP protein at the expected size of 96 kDa compared to the 29 kDa GFP alone (**Fig. S1C**).

Following successful generation of the NDC80-GFP transgenic line, the spatiotemporal profile of NDC80-GFP protein expression and location was examined during the parasite life cycle at the three asexual mitotic replicative stages (liver and blood schizogony in the vertebrate host and oocyst development (sporogony) in the mosquito vector) (**Fig. 1A**), the sexual mitotic stage (male gametogenesis) (**Fig. 1B**) and the meiotic stage (ookinete development) (**Fig. 1C**).

### Real-time live cell imaging using NDC80-GFP reveals kinetochores aggregate as discrete foci during schizogony

In the asexual blood stage, no NDC80-GFP fluorescence was observed in the intra-erythrocytic ring stage, a non-replicative G1 phase of the cell cycle (Arnot and Gull, 1998; Arnot et al., 2011; Doerig et al., 2000) (**Fig. 2A**). A faint but discrete single focus of NDC80-GFP adjacent to the nuclear DNA was observed in the early trophozoite, which became more intense as the trophozoite developed (**Fig. 2A**). The late trophozoite stage marks the transition into early S phase of the cell cycle, when DNA synthesis starts. The NDC80-GFP focus then split into two foci that migrated away from each other but remained attached to the nuclear DNA (stained using Hoechst) that then separated into two nuclear masses. This is consistent with the separation of sister chromatids (anaphase) and then the first nuclear division (telophase) that marks the onset of schizogony. These observations indicate that kinetochores are grouped together in a tight focus throughout these mitotic stages of nuclear replication. Similar fluorescence patterns were also observed in the NDC80-mCherry line (**Fig. S2**). NDC80-GFP also revealed the asynchronous nature of nuclear division during early schizogony, as displayed by two or more nuclei with single and double distinct NDC80 foci concurrently (e.g. **Fig. 2A**, Sch-E). As alternating repeated S/M phases following the division of individual nuclei continued, these NDC80-GFP foci were duplicated several times into multiple foci and nuclei. Further analysis of NDC80-GFP localization by super resolution microscopy confirmed the asynchronicity of nuclear division during blood stage schizogony (see **Fig. 2B, Fig. S3A, B, C and Supplementary videos SV1, SV2, SV3**). This stage of DNA replication and nuclear division concludes with cytokinesis to produce haploid daughter merozoites that egress from the erythrocyte (Arnot and Gull, 1998; Doerig et al., 2000). The short-lived, extracellular merozoite represents part of the G1 phase of the cell cycle (Arnot and Gull, 1998; Doerig et al., 2000). During schizogony in the pre-erythrocytic asexual stage in the liver, discrete fluorescent foci next to nuclear DNA were observed, which is similar to the pattern observed in blood stage schizogony (**Fig. S3D**).

**Fig. 2:**
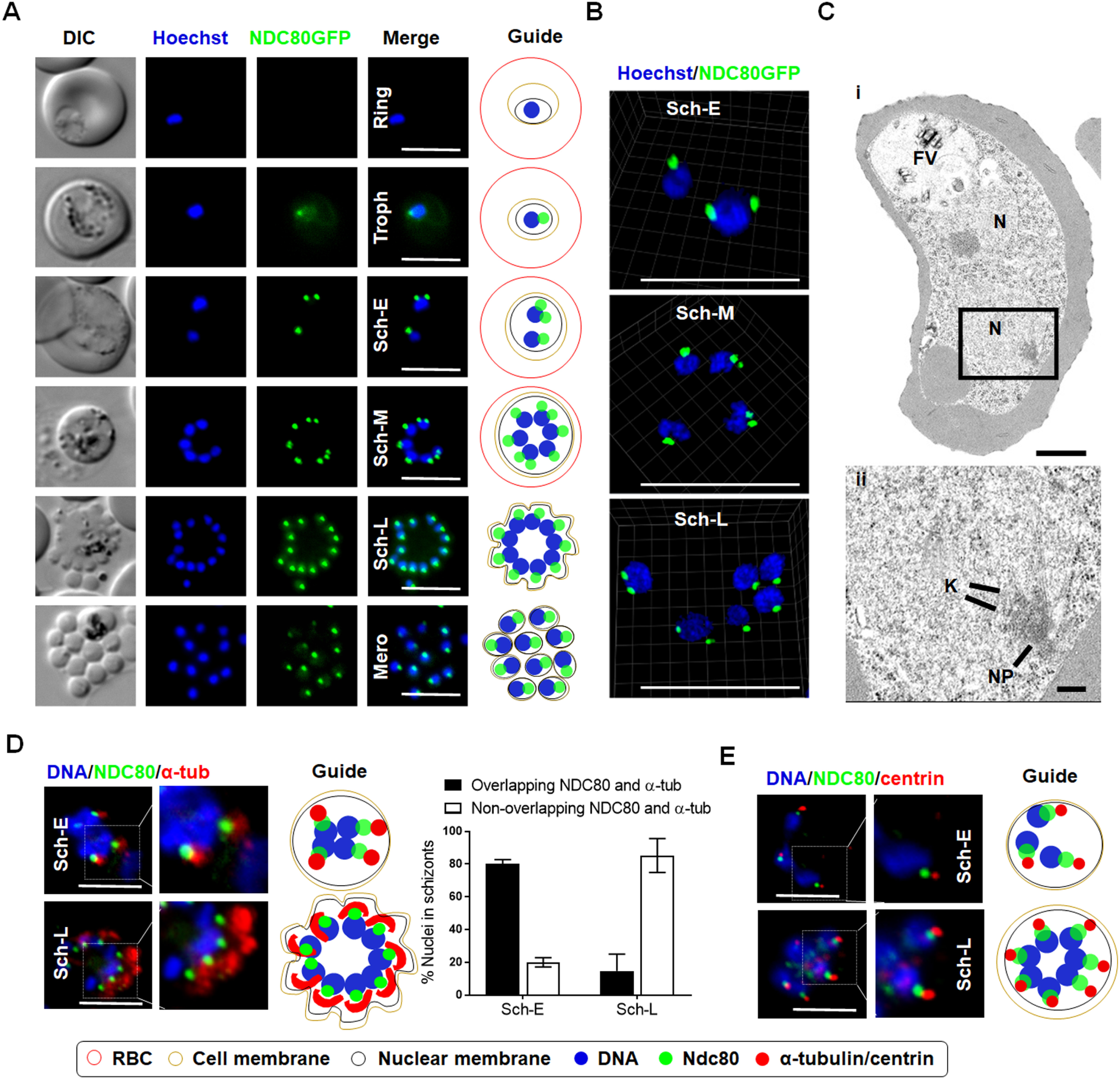
NDC80-GFP localization during endomitotic cell division in schizogony. **(A)** Live cell imaging of NDC80-GFP expression and location during asexual blood stage (DIC: Differential interference contrast, Hoechst: DNA, NDC80-GFP: GFP, Merge: Hoechst and GFP fluorescence, Troph: Trophozoite, Sch-E: Early Schizont, Sch-M: Mid Schizont Sch-L: Late Schizont, Mero: Merozoite, 100x magnification, Scale bar = 5 µm.) and schematic guide depicting NDC80 localization during various developmental stages in the bloodstream of the parasite life cycle. **(B)** Live cell super-resolution 3D imaging for NDC80-GFP localization during asynchronous blood stage mitotic division. Scale bar = 5 µm. (**C)** Electron microscopy imaging of an early schizont showing kinetochore localization. **(i)** Section through an early schizont within a red blood cell, showing two nuclei (N) and the food vacuole (FV). Scale bar = 1 µm. **(ii)** Enlargement of the enclosed area showing part of the nucleus in which a nuclear pole (NP), microtubules and attached kinetochores (K) can be seen. Scale bar = 100 nm. (**D)** Immunofluorescence fixed cell imaging and schematic guide of NDC80-GFP and co-localization with α-tubulin showing overlap during early schizont (**E**) Immunofluorescence fixed cell imaging and schematic guide of NDC80-GFP and co-localization with centrin (100x magnification). Scale bar = 5 µm.

To complement the live-imaging analysis, erythrocytic schizogony was examined in ultrastructural studies. In the multinucleated schizont during merozoite formation, it was possible to identify nuclear/spindle poles directly adjacent to the nuclear envelope with radiating microtubules and attached kinetochores on the inside of the nucleus **(Fig. 2C)**. These structures were not seen in the mature merozoite or the early intracellular ring stage. These observations are consistent with the live-imaging data.

### Immunofluorescence imaging shows the arrangement of NDC80 (kinetochore) with centrin (putative MTOC/SPB) and α-tubulin (spindles)

To determine the relative position of NDC80-GFP with other mitotic markers, especially at the spindle and/or putative MTOC/SPB, we used immunofluorescence-based co-localization assays with anti-GFP antibodies, anti-α-tubulin and anti-centrin, as markers for the spindle and putative MTOC/SPB, respectively. We observed that NDC80-GFP is located adjacent to alpha-tubulin showing some overlap between the NDC80 and alpha-tubulin signals in most early schizonts (Sch-E) but not in late schizonts (Sch-L) (**Fig. 2D).** Similarly, using anti-GFP with anti-centrin antibodies revealed that NDC80-GFP is located in close proximity to, but does not co-localize with, centrin (**Fig. 2E**)

### Rapid spindle dynamics of NDC80-GFP shows unusual kinetochore bridges during endoreduplication in male gametogenesis

Given the remarkable speed and organisation of nuclear replication and chromosome segregation during male gametogenesis, we investigated the live cell dynamics of NDC80-GFP throughout this 15 min process following gamete activation *in vitro*. The results are presented in **Fig. 3A**, and include time-lapse screenshots (**Fig. 3B and C, Supplementary videos SV4 and SV5.**). In non-activated male gametocytes a single diffuse and faint NDC80-GFP focus was present (**Fig S4A**), which intensified to a sharp single focal point 1 minute post-activation (mpa) (**Fig. 3A**). By 2 mpa, this focal point extended to form a bridge across one side of the nucleus, followed by the separation of the two halves of the bridge to produce two shorter linear rods that then contracted to two clear single foci by 3 mpa (**Fig. 3A-C)**. A schematic diagram for this process (1-3 mpa) is shown in **Fig. 3D.** This unusual linear arrangement of NDC80 shows that the distribution of kinetochores extends to approximately the full width of the nucleus, often arched around the nuclear margin, and maintains a consistent thickness of NDC80-GFP signal, suggesting that kinetochores are evenly spaced along this single linear element. This process was repeated twice, although non-synchronously, resulting in 8 discrete NDC80-GFP foci (**Fig. 3A**). To study the association of NDC80 with the spindle we used immunofluorescence-based co-localization assays with anti-GFP antibodies, and anti-α-tubulin. This showed clear co-localisation of NDC80 with the microtubule marker, both on the bridge-like structure and the foci for NDC80 (**Fig.3E**)

**Fig. 3:**
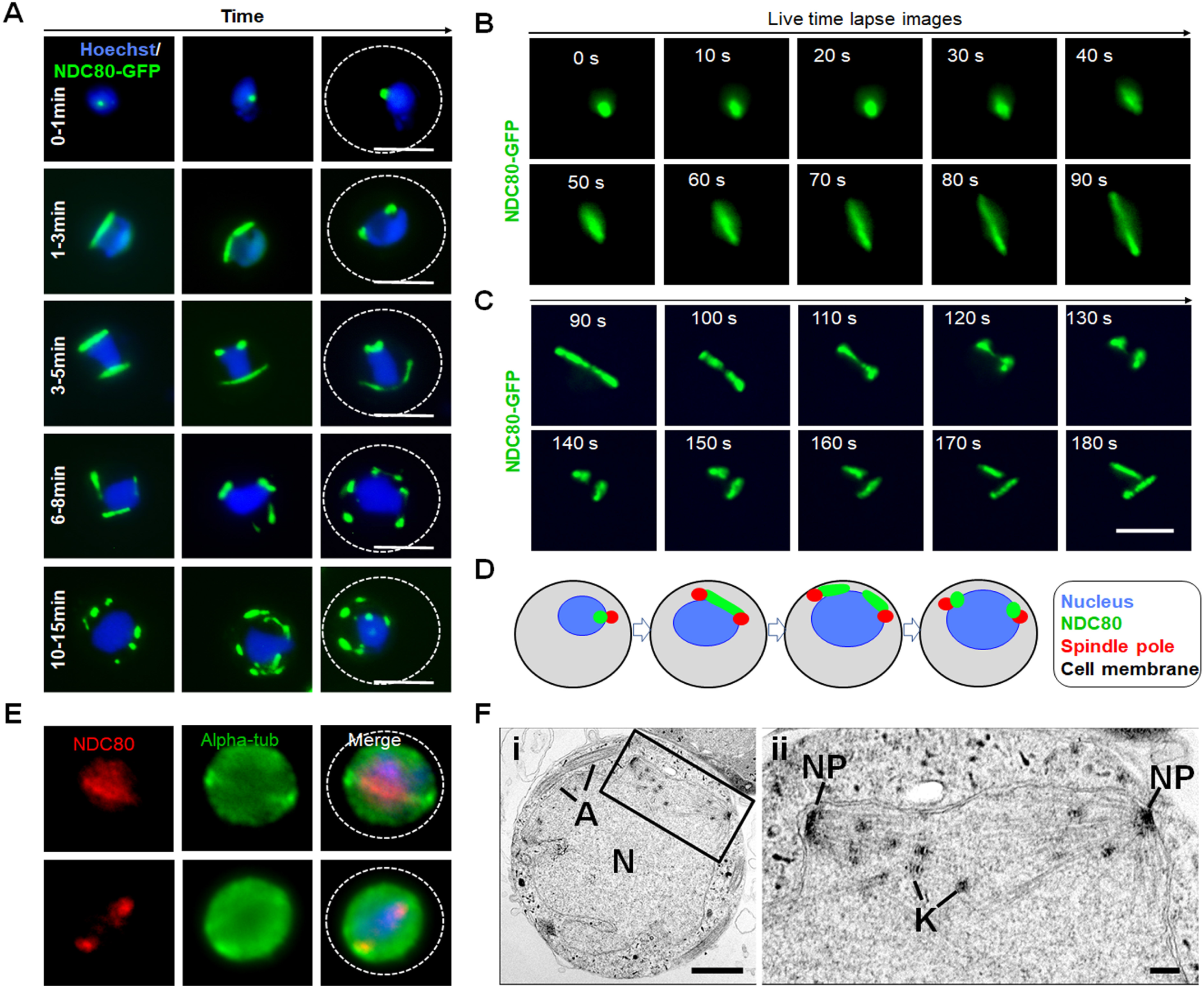
Temporal dynamics of NDC80-GFP during male gametogenesis. **(A)** Live cell imaging for NDC80-GFP expression and location during endoreduplicative mitotic division in male gametogenesis (100x magnification) **(B-C)** Time-lapse screenshots for NDC80-GFP localization during male gametogenesis. Scale bar = 5 µm. (**D)** Schematic representation showing dynamic localization of NDC80 during 1-3 min post activation of gametocyte (first round of nuclear division). **(E)** Indirect immunofluorescence assays showing co-localization of NDC80 (red) and α-tubulin (green) in male gametocytes activated for 1-3 min. **(F)** Electron microscopy on 8 min post-activation gametocytes for kinetochore localization. **(i)** Section through a mid-stage microgametocyte showing the large central nucleus (N) with axonemes (A) present in the peripheral cytoplasm. Scale bar = 1 µm. (**ii)** Enlargement of the enclosed area showing the details of an intranuclear spindle with microtubules with attach kinetochores (K) radiating from the nuclear poles (NP). Scale bar = 100nm.

Ultrastructural analysis of the nucleus during male gametogenesis within 8 mpa showed typical nuclear spindles with microtubules radiating from the nuclear poles, to which attached kinetochores could be identified dispersed along the length of the spindle (**Fig. 3F**). The spindles with attached kinetochores were located on the inner side of the nuclear membrane, (**Fig. 3F**). TEM images showed that these kinetochores are dispersed along the length of the mitotic spindle from one spindle pole to the other (similar to the bridge observed in the fluorescence microscopy). These observations are consistent with the fluorescence microscopy, and this kinetochore configuration is different to that in canonical metaphase, in which there is a central metaphase plate perpendicular to the spindle axis.

Following exflagellation, the eight discrete foci associated with endoreduplication disappeared rapidly and no NDC80-GFP fluorescence was observed in flagellated motile microgametes (**Fig. S4A**). Furthermore, no NDC80-GFP fluorescence was observed in either non-activated or activated female gametocytes (**Fig. S4A**). To independently test for an association of NDC80 with microtubules, we examined the effects of an anti-tubulin inhibitor (taxol) on NDC80 organisation during male gametogenesis. Addition of taxol at 1 min post activation blocked the dynamic progression of NDC80 distribution in more than 80% of male gametocytes while DMSO-treated gametocytes showed normal mitotic progression and NDC80 distribution **(Fig. S4B).** This showed that NDC80 distribution and localization depend on spindle dynamics and can be blocked by taxol treatment, which binds tubulin and stabilises microtubules by preventing depolymerisation. To further investigate the location of NDC80 in relation to the position of the centromere, NDC80-mCherry and the centromere-localised condensin core subunit SMC4-GFP parasite lines were crossed and used for live cell imaging of both markers to establish their spatiotemporal relationship, as shown previously (Pandey et al., 2019). The location of both NDC80 and SMC4 was next to the nucleus, showing co-localization **(Fig. S4C)**. Similarly, the basal body/axoneme marker, kinesin-8B-mCherry line (Zeeshan et al., 2019) was crossed with the NDC80-GFP line and this showed that NDC80 is located away from the basal body (kinesin-8B) during the start of male gametogenesis and later when kinesin-8B arranges across the axonemes (**Fig. S4D).** To confirm the centromere-associated localisation of NDC80 in a genome-wide manner, we performed a ChIP-seq experiment for NDC80-GFP in activated gametocytes **(Fig. S4E).** We observed strong ChIP-seq peaks at the centromeres of all 14 chromosomes similar to what was shown previously (Pandey et al., 2019).

### NDC80-GFP shows unusual dynamics throughout the meiotic stages during zygote to ookinete differentiation

Meiosis in the malaria parasite occurs during zygote differentiation to ookinete. This process takes 24 h to complete and during this time the ploidy of the parasite increases from 2N to 4N. To examine the behaviour of kinetochores throughout this process, we investigated the spatiotemporal profile of NDC80-GFP during ookinete differentiation.

NDC80-GFP fluorescence was first observed 1 to 1.5 h post-fertilization, as a single faint but distinct focal point, which gradually increased in intensity over the period 2 to 3 h post-fertilization (**Fig. 4**). As in male gametogenesis, no nuclear division was observed while the NDC80-GFP focus enlarged and divided to form a pair of elongated rod-like features. During stages II to IV of ookinete development, these rods appeared to fragment into multiple foci, ultimately resolving as 4 discrete foci in the mature ookinete, which has a 4N genome, at 18 h post-fertilization (**Fig. 4A**). The ultrastructure of the mature ookinete clearly showed four kinetochore clusters representing the 4N, but already segregated, genome within an intact nuclear membrane, consistent with the live cell imaging (**Fig. 4B**).

**Fig. 4:**
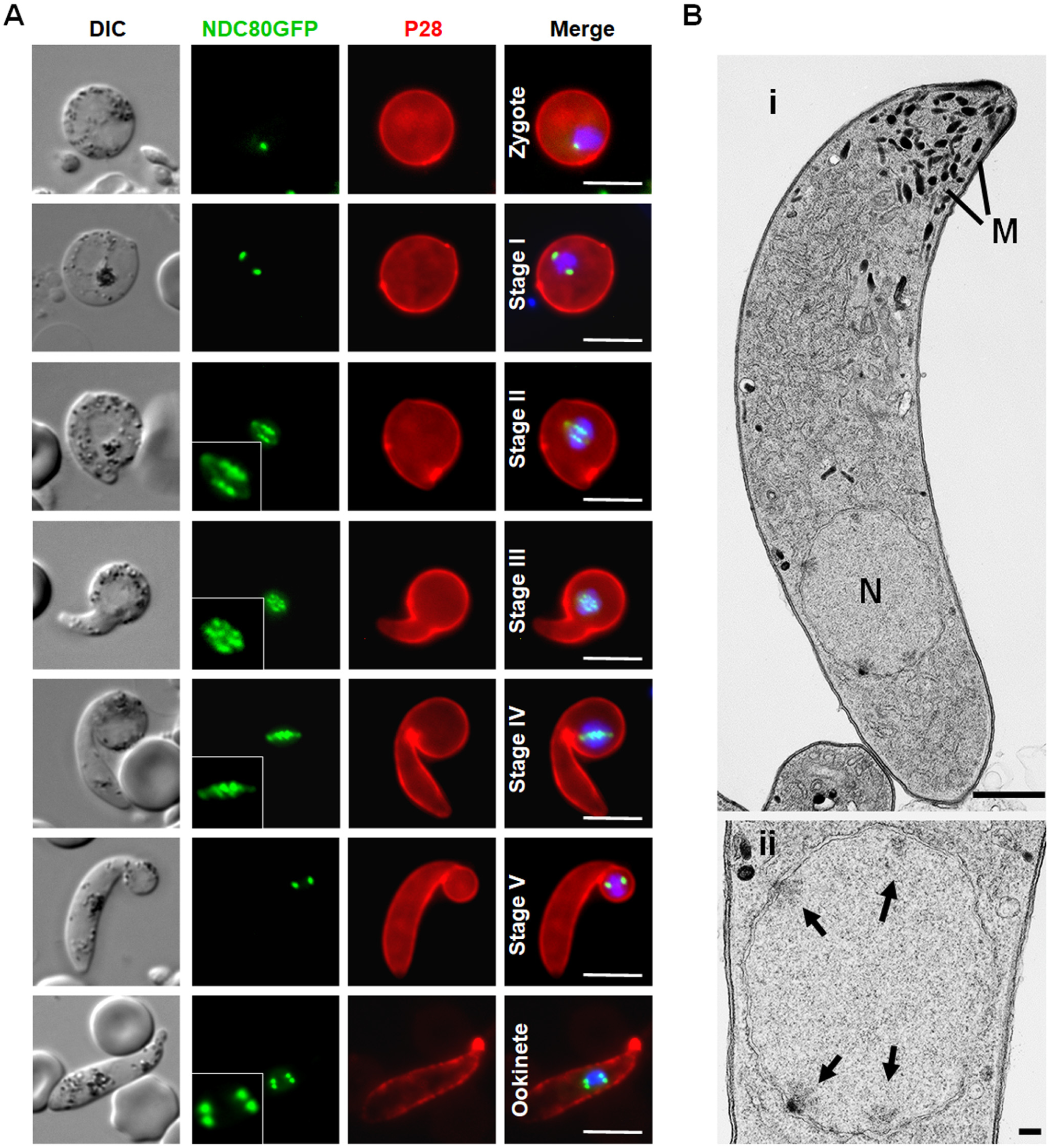
Spatiotemporal profile of NDC80-GFP expression during meiotic stages in the ookinete. (**A**) Live cell imaging of NDC80-GFP localization during various stages of ookinete development from zygote to mature ookinete in the mosquito gut (100x magnification). Merge: Hoechst (blue, DNA), GFP (green) and P28 (red, cell surface marker of activated female gamete, zygote and ookinete stages). Scale bar = 5 µm. (**B**) Ultrastructural analysis of kinetochore localization in a mature ookinete. (**i**) Longitudinal section through a mature ookinete showing the apical complex with several micronemes (M) and the more posterior nucleus (N). Scale bar = 1 µm. (**ii)** Enlargement of the nucleus showing the location of the four nuclear poles (arrows). Scale bar = 100 nm.

### NDC80-GFP is present as multiple foci during oocyst development and sporozoite formation

During oocyst development and sporozoite formation, live cell imaging revealed NDC80-GFP fluorescence at multiple foci adjacent to the nuclear DNA during various stages of oocyst development from 7 days post-infection (dpi) of the mosquito to 21 dpi, as well as a single focus in mature sporozoites (**Fig. 5A and B**). Ultrastructure analysis of oocyst development revealed an enlarged nucleus that formed a large multiple lobed structure with multiple nuclear poles/centriolar plaques/SPB/putative MTOC, followed by the formation of large numbers of sporozoites at the plasmalemma of the oocyst (**Fig. 5Bi**), as described previously (Ferguson et al., 2014; Schrevel et al., 1977). Detailed examination showed nuclear poles/centriolar plaques with kinetochores directed toward the developing sporozoites (**Fig. 5Bii**). The endomitotic process of sporozoite formation during sporogony resembles that of merozoite formation within host red cells and hence is similar to schizogony but with many more nuclei.

**Fig. 5:**
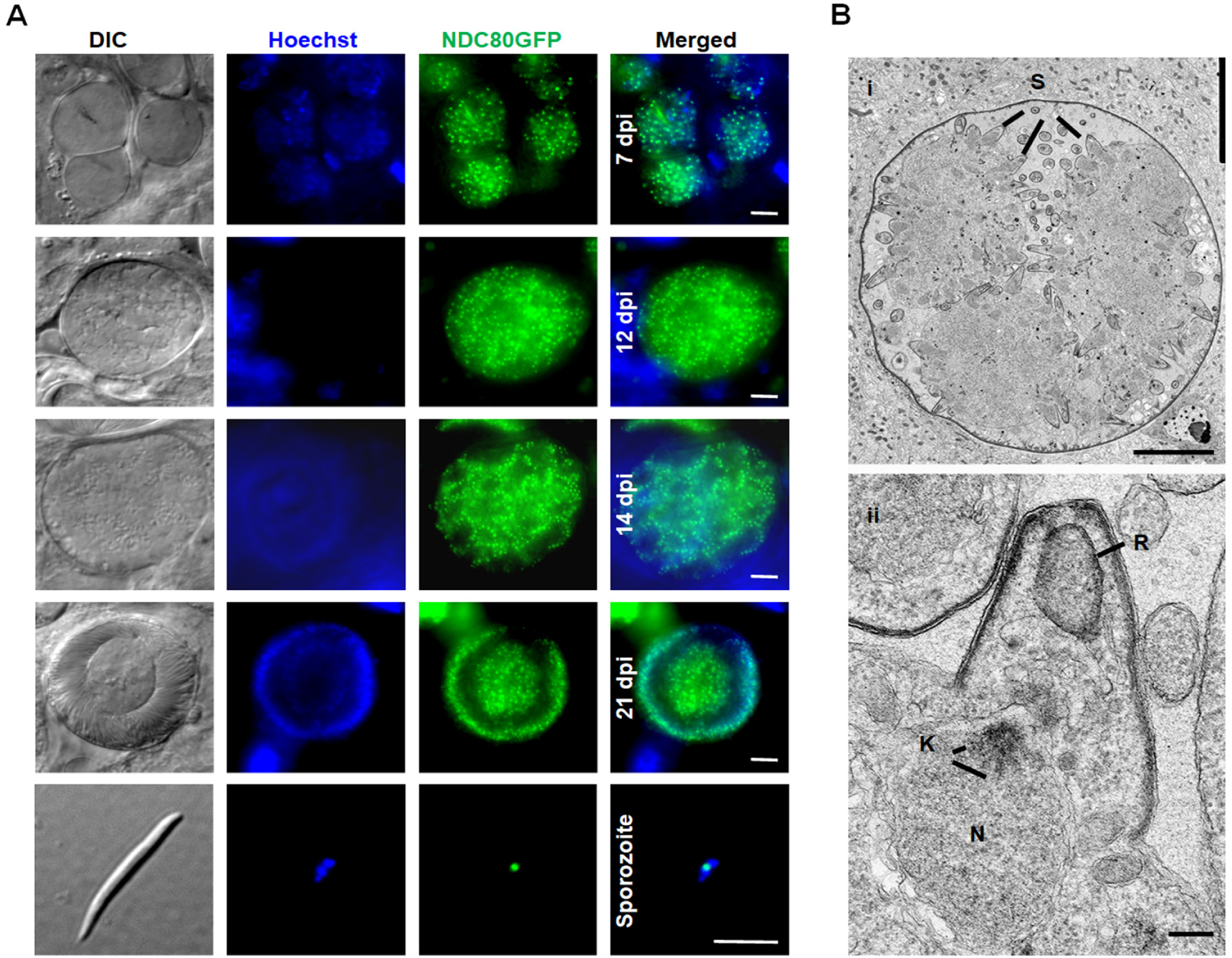
NDC80-GFP localisation during oocyst development and sporozoite formation. (**A**) Live cell imaging of NDC80-GFP in oocysts at 7, 12, 14 and 21 days post-infection (dpi) and a sporozoite. Panels: DIC (differential interference contrast), Hoechst (blue, DNA), NDC80-GFP (green, GFP), Merged: Hoechst (blue, DNA) and NDC80-GFP (green, GFP) Scale bar = 5 µm. **(B)** Electron microscopy analysis of kinetochore location in an oocyst 12 day post-infection. **(i)** Central section through a mid-stage oocyst showing the early stages of sporozoite formation (S) at the surface of the oocyst cytoplasm. Scale bar = 10 µm. **(ii)** Detail showing the early stage in sporozoite budding. Note the underlying nucleus (N) with the nuclear pole and attached kinetochores (K) directed toward the budding sporozoite. R – Rhoptry anlagen. Scale bar = 100 nm.

### Immunoprecipitation of NDC80-GFP recovers canonical members of the NDC80 complex and reveals a highly divergent SPC24-like candidate

Previous comparative genomics studies revealed evidence for the presence of three NDC80 complex members in Plasmodiidae: NDC80, NUF2 and SPC25, but did not identify any candidate SPC24 ortholog (van Hooff et al., 2017). In another apicomplexan lineage (*Cryptosporidium*), however, an SPC24 ortholog was identified, raising the question of whether SPC24 has been lost in Plasmodiidae or is an as-yet-unidentified highly divergent ortholog present. To determine the composition of the NDC80 complex in *P. berghei*, possibly including novel NDC80 interactors that might be responsible for the distinct kinetochore localisations in different life stages, we immunoprecipitated NDC80-GFP from lysates of schizonts following culture for 8-hours and from gametocyte lysates one minute after activation. Mass spectrometric analysis of these pulldowns identified NUF2 (PBANKA_0414300) as the main binding partner of NDC80, and we detected SPC25 (PBANKA_1358800) as part of a longer list of proteins identified with fewer unique peptides and recovered for the NDC80-GFP precipitate, but absent from the GPF-only control (**Fig. 6B, Table S2**). For further scrutiny of the list of candidate proteins, we assessed similarity in behaviour to NDC80 and/or SPC25 across the control and NDC80-GFP pulldown experiments using both principal component analysis (PCA), and Spearman rank correlation. We selected candidate proteins that showed similar variance to SPC25 based on the first two components of the PCA analysis, and those that had a correlation value of R>0.7 for both NDC80 and SPC25 (**Fig S5**). Of the resulting list, most proteins have functions in transcription/RNA-related processes, and one is the extracellular protein casein kinase 1 (PBANKA_0912100)(Dorin-Semblat et al., 2015), suggesting likely non-specific association within the cell lysate. However, one protein stood out: PBANKA_1442300, a protein with a long coiled-coil region and a predicted C-terminal globular domain, which suggested that it might be a SPC24 ortholog (**Fig. 6B**).

**Fig. 6:**
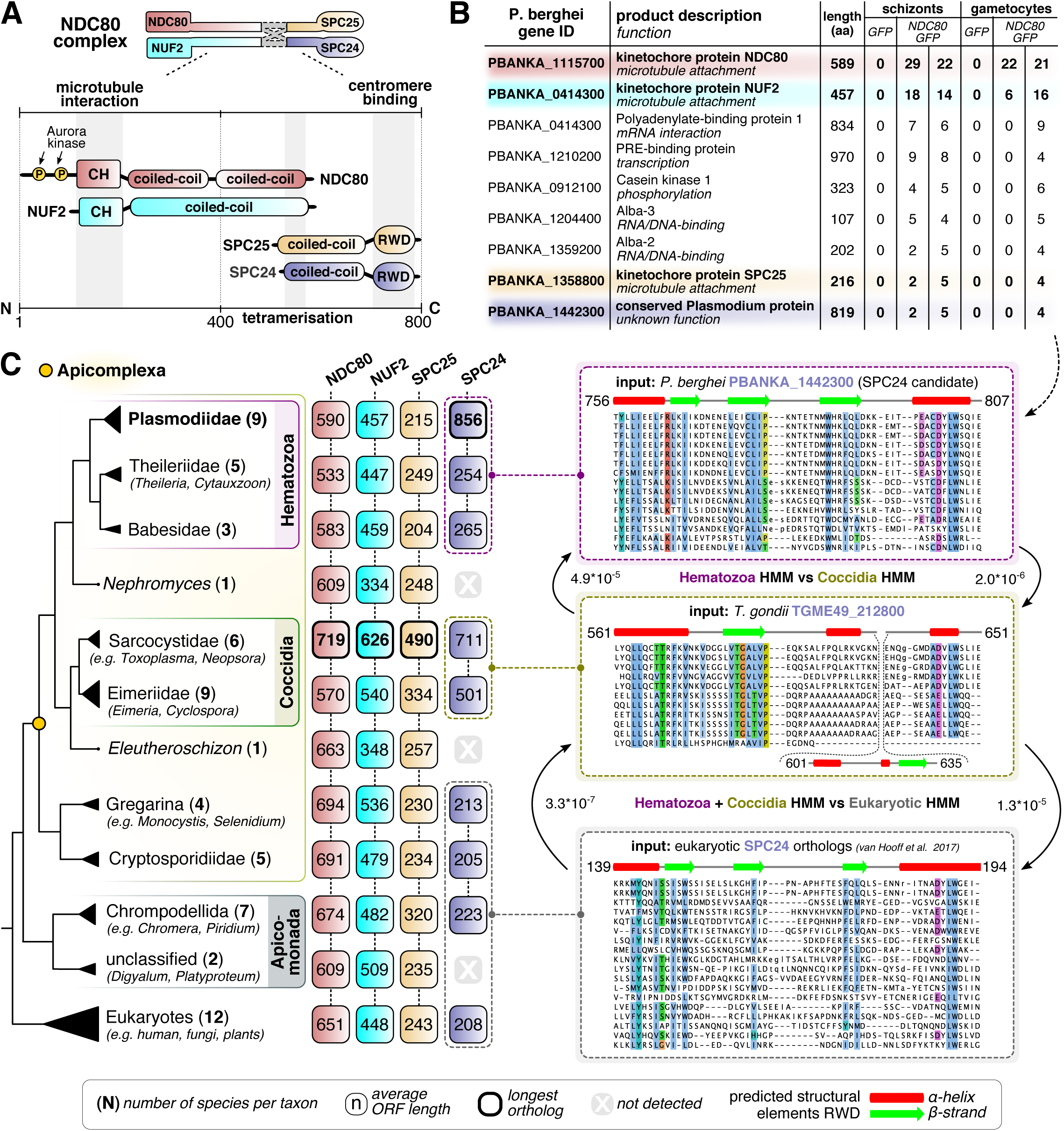
The NDC80 complex is conserved in *Plasmodium spp.* and most Apicomplexa. **(A)** Domain composition of the four subunits of the NDC80 complex tetramer as commonly found in model eukaryotes. N and C denote the amino and carboxy termini of the proteins. Numbers indicate the length in amino acids. **(B)** List of candidate proteins identified by mass spectrometry analysis of anti-GFP immunoprecipitants from lysates of WT-GFP and NDC80-GFP schizonts (after 8 hr in culture) and gametocytes (activated for 1 min) (see **Fig. S5**). Numbers are the total number of peptides identified for each protein. We identify the previously unannotated PBANKA_1442300 gene as coding for a candidate SPC24 ortholog in *P. berghei* (see panel C). Colours correspond to panel A and indicate each of the four subunits of NDC80 complex. **(C)** *On the left:* presence/absence matrix of NDC80 complex subunits in a large set of (newly sequenced) apicomplexan parasites, various Apicomplexa-affiliated lineages (outgroup e.g. chrompodellids) and a subset of eukaryotes. (see **Table S4** for presence/absence table, and **Sequence File S1** for sequences of NDC80 complex orthologs). Numbers indicate the average length of the orthologs in each collapsed clade represented in the phylogenetic tree. Note the consistently longer orthologs in Sarcocystidae (including *T. gondii*) and the expanded SPC24 ortholog in Plasmodiidae. *On the right:* workflow for the discovery of SPC24 candidate orthologs in Hematozoa and Coccidia. Alignments represent RWD domains of putative SPC24 orthologs found using seed sequences (top: e.g. TGME49_212800). See **Fig. S6** for full alignments of RWD domains. The panels represent Hidden Markov Models (HMMs) of SPC24-like RWD domains, which were found to be significantly similar (see E-values) using HMM-vs-HMM search algorithm HHsearch (see methods).

To test whether this candidate was a genuine SPC24 ortholog, we first queried available databases for PBANKA_1442300 homologs using conventional sequence similarity detection approaches, but detected none outside of hematozoan lineages, consistent with these approaches having failed to discover Plasmodiiae SPC24 candidates in the past (Plowman et al., 2019; van Hooff et al., 2017). Therefore, we employed more sophisticated protein modelling approaches to discover PBANKA_1442300 orthologs. We constructed a large sequence database consisting of 64 apicomplexan and other eukaryotic genomes and transcriptomes (see **Table S3** for sources of the sequence database) and generated Hidden Markov Models (HMM) of automatically defined homologous groups of sequences (see methods). We then compared these models with HMM profiles of PBANKA_1442300-like homologs and bona fide eukaryotic SPC24 orthologs found in our dataset. This multi-step approach yielded candidate apicomplexan SPC24 orthologs (**Fig. 6C**). Hematozoan homologs (including PBANKA_1442300) were significantly similar (E > 10^−5^) to the previously unannotated group of coccidian homologous sequences (including the *Toxoplasma gondii* gene TGME49_212800). The merged HMM profile of the C-terminal globular domain of these two groups, in turn, was significantly similar to that of eukaryote-wide SPC24 orthologs (E > 10^−5^) (**Fig. 6C, Fig. S6**). These analyses provide strong credence to the idea that PBANK_1442300- and TGME49_212800-like sequences are divergent but bona fide homologs of SPC24. Given that no other SPC24 candidates were found in these taxa, and the proteomic evidence of association of PBANKA_1442300 with SPC25, NDC80 and NUF2, it is very likely that PBANK_1442300 and TGME49_212800 are genuine SPC24 functional orthologs (**Fig. 6C, Sequence File S1, Table S4**).

## Discussion

Cellular proliferation in eukaryotes requires chromosome replication and precise segregation, followed by cell division, to ensure that daughter cells have identical copies of the genome. This happens through assembly of a spindle to which the centromeric region of chromosomes is attached through the kinetochore. Although the organisation of spindle microtubules, the molecular composition of kinetochores, and the modes of spindle pole separation vary extensively among eukaryotes (Akiyoshi and Gull, 2013; Drechsler and McAinsh, 2012; van Hooff et al., 2017), the microtubule-binding subunit NDC80 is conserved across most eukaryotes including apicomplexan parasites such as *Toxoplasma gondii* and *Plasmodium spp.* (Akiyoshi and Gull, 2013; Farrell and Gubbels, 2014; van Hooff et al., 2017). In addition, since chromosomes only bear one kinetochore, the outer-kinetochore subunit NDC80 is an excellent tool to start probing the rather surprising chromosome dynamics during the different stages of the life cycle in *P. berghei*.

In this study we have sought to understand the assembly and dynamics of the mitotic machinery during the diverse modes of nuclear division in *Plasmodium* using the kinetochore-protein NDC80 as a marker for chromosome attachment to the mitotic spindle. For this we generated a transgenic parasite line to express endogenous C-terminal GFP-labelled NDC80, a protein which faithfully replicates kinetochore location and function during all diverse mitotic and meiotic stages of the life cycle. Live-cell imaging of the fluorescent protein, complemented with ultrastructural studies by electron microscopy, revealed a subcellular location of NDC80-GFP at discrete foci adjacent to nuclear DNA in all replicative stages of the *P. berghei* life cycle. The distribution and dynamic spatiotemporal profile corresponded to the replication of chromosomes during the atypical mitotic and meiotic processes of DNA replication in this organism. Non-replicating stages, including the intraerythrocytic ring stage, the extracellular mature merozoite, the non-activated female gametocyte and the motile male gamete show no evidence of kinetochore assembly as no NDC80-GFP expression was observed. Using a combination of GFP-pulldown and a sensitive homology detection workflow we identified all four components of the NDC80 complex throughout apicomplexans, including a likely SPC24 ortholog candidate in Plasmodiidae.

The subcellular localization data for NDC80-GFP revealed a discreet single focus adjacent to the haploid nuclear genome, which presumably contains the centromeres of all 14 chromosomes. Such clustering of centromeric regions has been demonstrated in yeast (Richmond et al., 2013), human cells (Solovei et al., 2004) and *Toxoplasma* (Farrell and Gubbels, 2014). It is thought to be important for genome integrity, but the exact reason for it is not well understood. In the stages of the life cycle where this NDC80 clustering is not detected, perhaps the 14 kinetochores are not fully assembled and therefore NDC80-GFP expression is not detectable (Hoeijmakers et al., 2012). This is consistent with the idea that clustering of centromeres in *Plasmodium falciparum* occurs only prior to the onset of chromosome segregation (Hoeijmakers et al., 2012). It is of interest that following fertilization there is a single focus despite the genome being diploid, this may well reflect the tight pairing of sister chromatids allowing recombination to occur at this stage.

Although our data showed a clustered location of NDC80, its actual role in chromosome clustering is not known. Previous studies on yeast and *Toxoplasma* showed no role of NDC80 in clustering and suggested a sole role in attachment to spindle MTs during chromosome segregation. A single MT binds each kinetochore in budding yeast (Westermann et al., 2007); whereas in *Toxoplasma* a maximum of only 11 MTs were detected despite the fact that there are 13 chromosomes (Bunnik et al., 2019; Farrell and Gubbels, 2014; Swedlow et al., 2002). Within the closely related Coccidian parasites, a number of variations in the details of the process of asexual division has been described, relating to timing and number of genome and nuclear divisions (Ferguson et al., 2008) The coccidian parasite *Sarcocystis neurona*, which divides by endopolygeny forms a polyploid nucleus culminating in 64 haploid daughter cells (Farrell and Gubbels, 2014; Vaishnava et al., 2005). During this process, intranuclear spindle poles are retained throughout the cell cycle, which suggests constant attachment of chromosomes to spindle MTs via kinetochores to ensure genome integrity throughout *Sarcocystis* cell division (Farrell and Gubbels, 2014; Vaishnava et al., 2005). In contrast, *Plasmodium* (a hemosporidian), undergoes classical schizogony with a variable number of cycles of genome replication and nuclear division resulting a multinucleated cell (Arnot et al., 2011). This is similar to what is seen in the coccidian parasites *Eimeria* spp. and *Toxoplasma*, with daughter cell formation being associated with the final nuclear division. This fact is well demonstrated by the NDC80 localization in this study, which shows 1 or 2 NDC80-GFP foci per nucleus. The asynchronous nature of the division during these stages is shown by nuclei having either one or two NDC80-GFP foci (and intermediate forms) within the same cell. We made similar observations in our previous study of the SPB/MTOC/centriole plaque marker for centrin, CEN-4 (Roques et al., 2019). These studies also suggest that the kinetochore and centrosome/centriolar plaque duplicate before nuclear division starts; therefore the duplication of NDC80 and CEN-4 sets the stage for mitosis at each round of nuclear division in *Plasmodium*, as in *Toxoplasma* (Suvorova et al., 2015). In contrast, during male gametogenesis the genome size increases to 8N, and this corresponds to the formation of 8 distinct NDC80-GFP foci, before nuclear division, with asynchronous chromosome replication and segregation. Most notable is the presence of unique kinetochore bridges or rod-like structures during genome replication and chromosome segregation. It appears that two hemi-spindles associated with the kinetochores are joined together to form a mitotic spindle at the earliest stages of endoreduplication during male gametogenesis. This is then followed by kinetochore movement to opposite poles once all the duplicated chromosomes are segregated. The mitotic process of duplication proceeds in the absence of nuclear division (karyokinesis), which results in the very atypical kinetochore dynamics that are consistent with the live cell imaging data.

Similarly, during meiosis in ookinete development the genome size increases to 4N, represented by bridge-like kinetochore structures during replicative stages and at the end, resulting in four distinct NDC80-GFP foci suggesting kinetochore clustering to facilitate chromosome segregation, although no nuclear division takes place.

During sporogony multiple lobes are formed and the intranuclear spindle may be formed during multiple nuclear divisions, as revealed by ultrastructural studies during sporozoite formation (Schrevel et al., 1977). Further ultrastructure analyses identified typical nuclear spindles with attached kinetochores, radiating from the nuclear poles located within an intact nuclear membrane during schizogony, male gametogenesis, ookinete development and sporogony. Our data are consistent with a previous report, in which kinetochore localization was revealed using ultrastructural studies of *P. berghei* sporogony, and the duplication of hemi-spindles during replication was suggested (Schrevel et al., 1977). Overall, based on all these results, consistent kinetochore clustering occurs within Apicomplexa.

NDC80 is a major constituent of kinetochores and is highly conserved among eukaryotes including *Plasmodium.* However, many of the molecular details of kinetochore architecture and function in *Plasmodium* remain to be explored. A recent comparative evolutionary analysis suggested a distinct kinetochore network in many eukaryotes including *P. falciparum* and other alveolates, with many of the highly conserved kinetochore complex proteins being absent in *Plasmodium* (van Hooff et al., 2017). Of 70 conserved kinetochore proteins only eleven were found to be encoded in the *P. falciparum* genome. These eleven proteins include NDC80, NUF2, CENP-C/-A/-E, SPC25 and others that are highly conserved across 90 eukaryotic species, but genes for many other conserved proteins like SPC24, MAD-1/-2 and MIS12 were found to be absent (van Hooff et al., 2017). Recent studies have shown the presence of unconventional kinetochore proteins in kinetoplastids (Akiyoshi and Gull, 2013; D’Archivio and Wickstead, 2017), suggesting that different kinetochore architectures are possible. Previous reports have shown the association of CENP-A and CENP-C with centromeres in *P. falciparum* (Verma and Surolia, 2013; Verma and Surolia, 2014). Using a combination of GFP-pulldown and a sensitive homology detection workflow we identified all four components of the NDC80 complex throughout apicomplexans, including a highly divergent SPC24 ortholog candidate in Plasmodiidae (PBANKA_1442300). Previous studies in *Toxoplasma gondii* only identified NDC80 and NUF2 (Farrell and Gubbels, 2014), but here we predict the presence of SPC25 (TGME49_232400) and a SPC24-like homolog (TGME49_212800) in this organism. We favour the interpretation of PBANKA_1442300 being a bona fide SPC24 ortholog for two reasons: (1) it was detected as a putative NDC80-GFP binding partner, and (2) our sequence analyses indicate it has a C-terminal RWD-like domain most similar to that of SPC24. It seems unlikely that the ancestor of hematozoa and coccidia would have duplicated the gene and then discarded the functional kinetochore SPC24 ortholog. Further proteomic and co-localisation studies of SPC25 and SPC24 with each other as well as other kinetochore markers (for example, NUF2 and CENP-C) will be needed to confirm that the candidate SPC24 ortholog is a true kinetochore protein in *Plasmodium*. Our approach has illustrated that conspicuous absences of subunits of highly conserved and essential complexes, such as the obligate tetrameric NDC80 complex, are to be treated with caution, and additional sensitive homology searches using HMM-HMM comparison should be employed to more thoroughly test for homolog presences or absences. As such, patterns of extensive loss have been observed in eukaryotic parasites before, and we expect that systematic searches for conspicuous absences of subunits of specific complexes will yield a large number of highly divergent homologs in Apicomplexa.

NDC80 and NUF2 are the most conserved subunits of the NDC80 complex, whereas the SPC24-SPC25 dimer that interacts with the centromere is highly divergent both in sequence and in length. Strikingly, the candidate SPC24 orthologs in Plasmodiidae and Coccidia are 3 to 4 times longer than those of other eukaryotes (**Fig. 6C**), and their RWD domains are also relatively divergent. What this means exactly is unclear, but we speculate that the larger NDC80 complex present in some but not all Apicomplexa engages centromere-proximal kinetochore proteins in a different way than in other eukaryotes, and possibly through direct interaction with CENP-A and CENP-C. This notion is supported by the apparent absence of the canonical interaction partners of SPC24 and SPC25, namely CENP-T and the MIS12 complex (van Hooff et al., 2017), and the extended length of all NDC80 complex members within coccidia (∼1.2 - 3 times longer). In Plasmodiidae only the SPC24 candidate ortholog is extended (∼3 to 4 times longer), but in Coccidia, SPC25 is also twice as long as canonical SPC25s, potentially resulting in a larger and longer NDC80 complex. The extended length of SPC24 candidate orthologs results largely from N-terminal extensions of the coiled-coil region (Coccidia and Plasmodiidae), and/or large insertions into the loop of the RWD domain (Coccidia). We envision that these extensions may provide additional binding sites for novel interactors important in kinetochore clustering and/or the remarkable ‘bridge/rod’ phenotype we have observed in this study. The SPC24 N-terminal coiled-coil extension may also have additional interactions with the heterodimeric coiled-coils of the NDC80 and NUF2 subunits, providing extra rigidity to the tetrameric NDC80 superstructure.

In summary, this study demonstrates the dynamic expression and location of NDC80 during the different proliferative stages of the malaria parasite and reveals both the disassembly and reassembly, as well as clustering, of kinetochores. It also shows the asynchronous closed mitotic division of *Plasmodium* during schizogony and sporogony and provides novel insights into the chromosome segregation in male gametogenesis and in various stages of meiosis during zygote differentiation to ookinetes. ChIP-seq and colocalisation studies clearly showed the centromeric location of NDC80 in *Plasmodium*. The protein pulldown and bioinformatics studies revealed that NDC80 has the full complement of four subunits, though SPC24 is highly divergent compared to other eukaryotes. This analysis of NDC80 will also facilitate future studies of cell division and comparative analyses of chromosome dynamics in evolutionarily divergent eukaryotic cells.

## Material and Methods

### Ethical statement

All animal-related work performed at the University of Nottingham has undergone an ethical review process and been approved by the United Kingdom Home Office with the project license number 30/3248 and PDD2D5182. The work has been carried out in accordance with the United Kingdom ‘Animals (Scientific Procedures) Act 1986’ and was in compliance with ‘European Directive 86/609/EEC’ for the protection of animals used for experimental purposes. A combination of ketamine followed by antisedan was used for general anaesthesia and sodium pentobarbital was used for terminal anaesthesia. Proper care and efforts were made to minimise animal usage and suffering.

Six to eight-week-old female Tuck-Ordinary (TO) (Harlan) or CD1 outbred mice (Charles River) were used for all experiments.

### Generation of transgenic parasites

The transgenic lines for NDC80 (PBANKA_1115700) were created using single homologous recombination as shown in **Fig. S1**. The oligonucleotides used to generate transgenic lines are provided in **Supplementary Table S1**. For GFP tagging, a 1153bp region of *Ndc80* without the stop codon was inserted upstream of the *gfp* sequence in the p277 plasmid vector using KpnI and ApaI restriction sites as described previously (Tewari et al., 2010). The p277 vector contains the human *dhfr* cassette, conveying resistance to pyrimethamine. Before transfection, the sequence was linearised using EcoRV. The *P. berghei* ANKA line 2.34 was used for transfection by electroporation (Janse et al., 2006). Immediately, electroporated parasites were mixed with 100μl of reticulocyte-rich blood from a phenylhydrazine (6 mg/ml, Sigma) treated, naïve mouse and incubated at 37°C for 30 min before intraperitoneal injection. Pyrimethamine (70 mg/L, Sigma) was supplied in the drinking water from 1-day post-infection (dpi) to 4-dpi. Infected mice were monitored for 15 days and drug selection was repeated after passage to a second mouse. Integration PCR and western blot were performed to confirm successful generation of the transgenic line. For integration PCR, primer 1 (IntT259) and primer 2 (ol492) were used to confirm integration of the GFP targeting construct. Primer 1 and primer 3 (T2592) were used as a control. We also generated a mCherry-tagged NDC80 transgenic parasite line as shown in the schematic provided in **Fig. S1.**

### Western Blot

For western blotting, purified schizonts were lysed using lysis buffer (10 mM TrisHCl pH 7.5, 150 mM NaCl, 0.5 mM EDTA, 1% NP-40 and 1% Sarkosyl). The samples were boiled for 10 min after adding Laemmli sample buffer to the lysed cells. The sample was centrifuged at 13500 g for 5 min and electrophoresed on a 4–12% SDS-polyacrylamide gel. Subsequently, resolved proteins were transferred to nitrocellulose membrane (Amersham Biosciences) and immunoblotting was performed using the Western Breeze Chemiluminescence Anti-Rabbit kit (Invitrogen) and anti-GFP polyclonal antibody (Invitrogen) at a dilution of 1:1250, according to the manufacturer’s instructions.

### Localization of NDC80-GFP throughout the parasite life cycle

Live cell imaging of transgenic parasite lines was performed at different proliferative stages during the parasite life cycle (**Fig. 1**) as described previously (Roques et al., 2019; Saini et al., 2017) using a Zeiss AxioImager M2 microscope fitted with an AxioCam ICc1 digital camera (Carl Zeiss, Inc).

### Blood stage schizogony

Infected mouse blood provided asexual blood and gametocyte stages of the *P. berghei* life cycle. Schizont culture (RPMI 1640 containing 25 mM HEPES, 1:10 (v/v) fetal bovine serum and penicillin/streptomycin 1:100) at different time points was used to analyse various stages of asexual development from ring to merozoite. The periods used for analysis and imaging were 0 to 1 h for ring stage parasites, 2 to 4 h for trophozoites, 6 to 8 h for early and mid-stage schizonts, 9 to 11 h for late segmented schizonts, and 18 to 24 h for mature schizonts and released merozoites in schizont culture medium.

### Male gametocyte development

*In vitro* cultures were prepared to analyse non-activated gametocytes, activated gametocytes and male exflagellation. For *in vitro* exflagellation studies, gametocyte-infected blood was obtained from the tails of infected mice using a heparinised pipette tip. Gametocyte activation was performed by mixing 100µl of ookinete culture medium (RPMI 1640 containing 25 mM HEPES, 20% fetal bovine serum, 10 mM sodium bicarbonate, 50 µM xanthurenic acid at pH 7.6) with gametocyte-infected blood. To study different time points during microgametogenesis, gametocytes were purified using Nycodenz gradient (48%) and monitored at different time points to study mitotic division (male gametogenesis, 0 to 15 min post-activation (mpa).

### Ookinete development

To study ookinete development, gametocyte infected blood was incubated in ookinete medium for 24 hpa and various stages of zygote differentiation and ookinete development were monitored at different time points (0 min for nonactivated gametocytes, 30 min for activated gametocytes, 2 to 3 h for zygotes, 4 to 5 h for stage I, 5 to 6 h for stage II, 7 to 8 h for stage III, 8 to 10 h for stage IV, 11 to 14h for stage V, and 18 to 24 h for mature ookinetes post activation in ookinete medium).

### Oocyst and sporozoite development

For mosquito transmission stages and bite back experiments, triplicate sets of 30 to 50 *Anopheles stephensi* mosquitoes were used. The mosquito guts were analysed on different days post-infection (dpi): 7 dpi, 12 dpi, 14 dpi and 21 dpi to check expression and localization of NDC80GFP during oocyst development and sporozoite formation.

### Schizogony in liver stages

To study localization of NDC80-GFP in *P. berghei* liver stages, 100,000 HeLa cells were seeded in glass-bottomed imaging dishes. Salivary glands of female *A. stephensi* mosquitoes infected with NDC80-GFP parasites were isolated and sporozoites were released using a pestle to disrupt salivary gland cells. The released sporozoites were pipetted gently onto the HeLa cells and incubated at 37 °C in 5% CO_2_ in air, in complete minimum Eagle’s medium containing 2.5 μg/ml amphotericin B. For initial infection, medium was changed at 3 h post-infection and thereafter once a day. To perform live cell imaging, Hoechst 33342 (Molecular Probes) was added (1 μg/ml) and imaging was done at 55 h post-infection using a Leica TCS SP8 confocal microscope with the HC PL APO 63x/1.40 oil objective and the Leica Application Suite X software.

### Indirect immunofluorescence assay

IFA studies were performed using poly-L-lysine coated slides on which schizonts had been previously fixed in 2% paraformaldehyde (PFA) in microtubule stabilising buffer (MTSB:10 mM MES, 150 mM NaCl, 5 mM EGTA, 5 mM MgCl_2_, 5 mM glucose) in 1X-PBS for 30 min at room temperature (RT) and smeared onto slides. The fixed cells were permeabilized using TBS containing 0.2% TritonX-100 for 5 min and washed three times with TBS before blocking. For blocking, 1 hour incubation was performed with TBS solution containing 3% BSA (w/v) and 10% goat serum (v/v). TBS containing 1% BSA and 1% goat serum was used to dilute the antibodies for the incubations. Anti-GFP rabbit antibody (Invitrogen) was used at 1:250 dilution, anti-alpha-tubulin mouse antibody (Sigma-Aldrich) was used at 1:1000 dilution, and anti-centrin mouse clone 20h5 antibody (Millipore) was used at 1:200 dilution; each was incubated for 1 hour at RT. Three washes were performed with TBS, then AlexaFluor 568 labelled anti-rabbit (red) and AlexaFluor 488 labelled anti-mouse (green) (Invitrogen) (1:1000 dilution) were used as secondary antibodies and incubated for 40 min at RT. A similar protocol was followed for gametocytes except the cells were fixed in 4% PFA in MTSB. Slides were mounted with Vectashield containing DAPI (blue) and sealed using nail polish. Images were captured as described for live imaging.

### Super resolution microscopy

A small volume (3 µl) of schizont culture was mixed with Hoechst dye and pipetted onto 2 % agarose pads (5×5 mm squares) at room temperature. After 3 min these agarose pads were placed onto glass bottom dishes with the cells facing towards glass surface (MatTek, P35G-1.5-20-C). Cells were scanned with an inverted microscope using Zeiss C-Apochromat 63×/1.2 W Korr M27 water immersion objective on a Zeiss Elyra PS.1 microscope, using the structured illumination microscopy (SIM) technique. The correction collar of the objective was set to 0.17 for optimum contrast. The following settings were used in SIM mode: lasers, 405 nm: 20%, 488 nm: 50%; exposure times 100 ms (Hoechst) and 25 ms (GFP); three grid rotations, five phases. The band pass filters BP 420-480 + LP 750 and BP 495-550 + LP 750 were used for the blue and green channels, respectively. Multiple focal planes (Z stacks) were recorded with 0.2 µm step size; later post-processing, a Z correction was done digitally on the 3D rendered images to reduce the effect of spherical aberration (reducing the elongated view in Z; a process previously tested with fluorescent beads). Images were processed and all focal planes were digitally merged into a single plane (Maximum intensity projection). The images recorded in multiple focal planes (Z-stack) were 3D rendered into virtual models and exported as images and movies (see supplementary material). Processing and export of images and videos were done by Zeiss Zen 2012 Black edition, Service Pack 5 and Zeiss Zen 2.1 Blue edition.

### Chromatin Immunoprecipitation sequencing analysis (ChIP-seq)

For the ChIP-seq analysis, libraries were prepared from crosslinked cells (using 1% formaldehyde). The crosslinked parasite pellets were resuspended in 1 mL of nuclear extraction buffer (10 mM HEPES, 10 mM KCl, 0.1 mM EDTA, 0.1 mM EGTA, 1 mM DTT, 0.5 mM AEBSF, 1X protease inhibitor tablet), post 30 min incubation on ice, 0.25% Igepal-CA-630 was added and homogenized by passing through a 26G x ½ needle. The nuclear pellet extracted through 5000 rpm centrifugation, was resuspended in 130 µl of shearing buffer (0.1% SDS, 1 mM EDTA, 10 mM Tris-HCl pH 7.5, 1X protease inhibitor tablet), and transferred to a 130 µl Covaris sonication microtube. The sample was then sonicated using a Covaris S220 Ultrasonicator for 10 min for schizont samples and 6 min for gametocyte samples (Duty cycle: 5%, Intensity peak power: 140, Cycles per burst: 200, Bath temperature: 6°C). The sample was transferred to ChIP dilution buffer (30 mM Tris-HCl pH 8, 3 mM EDTA, 0.1% SDS, 30 mM NaCl, 1.8% Triton X-100, 1X protease inhibitor tablet, 1X phosphatase inhibitor tablet) and centrifuged for 10 min at 13,000 rpm at 4°C, retaining the supernatant. For each sample, 13 μl of protein A agarose/salmon sperm DNA beads were washed three times with 500 µl ChIP dilution buffer (without inhibitors) by centrifuging for 1 min at 1000 rpm at room temperature, then buffer was removed. For pre-clearing, the diluted chromatin samples were added to the beads and incubated for 1 hour at 4°C with rotation, then pelleted by centrifugation for 1 min at 1000 rpm. Supernatant was removed into a LoBind tube carefully so as not to remove any beads and 2 µg of anti-GFP antibody (ab290, anti-rabbit) were added to the sample and incubated overnight at 4°C with rotation. Per sample, 25 µl of protein A agarose/salmon sperm DNA beads were washed with ChIP dilution buffer (no inhibitors), blocked with 1 mg/mL BSA for 1 hour at 4°C, then washed three more times with buffer. 25 µl of washed and blocked beads were added to the sample and incubated for 1 hour at 4°C with continuous mixing to collect the antibody/protein complex. Beads were pelleted by centrifugation for 1 min at 1000 rpm at 4°C. The bead/antibody/protein complex was then washed with rotation using 1 mL of each buffers twice; low salt immune complex wash buffer (1% SDS, 1% Triton X-100, 2 mM EDTA, 20 mM Tris-HCl pH 8, 150 mM NaCl), high salt immune complex wash buffer (1% SDS, 1% Triton X-100, 2 mM EDTA, 20 mM Tris-HCl pH 8, 500 mM NaCl), high salt immune complex wash buffer (1% SDS, 1% Triton X-100, 2 mM EDTA, 20 mM Tris-HCl pH 8, 500 mM NaCl), TE wash buffer (10 mM Tris-HCl pH 8, 1 mM EDTA) and eluted from antibody by adding 250 μl of freshly prepared elution buffer (1% SDS, 0.1 M sodium bicarbonate). We added 5 M NaCl to the elution and cross-linking was reversed by heating at 45°C overnight followed by addition of 15 μl of 20 mg/mL RNAase A with 30 min incubation at 37°C. After this, 10 μl 0.5 M EDTA, 20 μl 1 M Tris-HCl pH 7.5, and 2 μl 20 mg/mL proteinase K were added to the elution and incubated for 2 hours at 45°C. DNA was recovered by phenol/chloroform extraction and ethanol precipitation, using a phenol/chloroform/isoamyl alcohol (25:24:1) mixture twice and chloroform once, then adding 1/10 volume of 3 M sodium acetate pH 5.2, 2 volumes of 100% ethanol, and 1/1000 volume of 20 mg/mL glycogen. Precipitation was allowed to occur overnight at −20°C. Samples were centrifuged at 13,000 rpm for 30 min at 4°C, then washed with fresh 80% ethanol, and centrifuged again for 15 min with the same settings. Pellet was air-dried and resuspended in 50 μl nuclease-free water. DNA was purified using Agencourt AMPure XP beads. Libraries were prepared using the KAPA Library Preparation Kit (KAPA Biosystems), and were amplified for a total of 12 PCR cycles (15 s at 98°C, 30 s at 55°C, 30 s at 62°C) using the KAPA HiFi HotStart Ready Mix (KAPA Biosystems). Libraries were sequenced using the NovaSeq 6000 System (Illumina), producing 100-bp reads.

FastQC (https://www.bioinformatics.babraham.ac.uk/projects/fastqc/), was used to analyze raw read quality. Any adapter sequences were removed using Trimmomatic (http://www.usadellab.org/cms/?page=trimmomatic). Bases with Phred quality scores below 25 were trimmed using Sickle (https://github.com/najoshi/sickle). The resulting reads were mapped against the *P. berghei* ANKA genome (v36) using Bowtie2 (version 2.3.4.1) or HISAT2 (version 2-2.1.0), using default parameters. Reads with a mapping quality score of 10 or higher were retained using Samtools (http://samtools.sourceforge.net/), and PCR duplicates were removed by PicardTools MarkDuplicates (Broad Institute). For Chip-seq analysis, raw read counts were determined to obtain the read coverage per nucleotide. Genome browser tracks were generated and viewed using the Integrative Genomic Viewer (IGV) (Broad Institute). Proposed centromeric locations were obtained from Iwanaga and colleagues (Iwanaga et al., 2010).

### Inhibitor studies

Gametocytes were purified as above and treated with 1 µM taxol (Paclitaxel, T7402; Sigma) at 1 min post activation (mpa) and then fixed with 4% Paraformaldehyde (PFA) at 8 mpa. DMSO was used as a control treatment. These fixed gametocytes were then examined on a Zeiss Axio Imager M2 microscope fitted with an AxioCam ICc1 digital camera (Carl Zeiss, Inc).

### Electron microscopy

Samples for different mitotic stages of parasite development including schizonts (24 hours in culture), activated male gametocytes (8 min post-activation), infected mosquito guts (12 to 14 days post infection) and the meiotic stage from the mature ookinete (24hours post-activation) were fixed in 4% glutaraldehyde in 0.1 M phosphate buffer and processed for electron microscopy as previously described (Ferguson et al., 2005). Briefly, samples were post-fixed in osmium tetroxide, treated in bloc with uranyl acetate, dehydrated and embedded in Spurr’s epoxy resin. Thin sections were stained with uranyl acetate and lead citrate prior to examination in a JEOL1200EX electron microscope (Jeol UK Ltd).

### Protein pulldown, Immunoprecipitation and Mass Spectrometry

Schizonts, following 8 hours *in vitro* culture, and male gametocytes 1 min post activation of NDC80-GFP parasites were used to prepare cell lysates. Purified parasite pellets were crosslinked using formaldehyde (10 min incubation with 1% formaldehyde in PBS), followed by 5 min incubation in 0.125M glycine solution and 3 washes with phosphate buffered saline (PBS, pH 7.5). Immunoprecipitation was performed using crosslinked protein lysate and a GFP-Trap^®^_A Kit (Chromotek) following the manufacturer’s instructions. Proteins bound to the GFP-Trap^®^_A beads were digested using trypsin and the peptides were analysed by LC-MS/MS. Mascot (http://www.matrixscience.com/) and MaxQuant (https://www.maxquant.org/) search engines were used for mass spectrometry data analysis. The PlasmoDB database was used for protein annotation. Peptide and proteins having minimum threshold of 95% were used for further proteomic analysis. The mass spectrometry proteomics data have been deposited to the ProteomeXchange Consortium via the PRIDE partner repository with the dataset identifier PXD017619 and 10.6019/PXD017619.

To assess covariance among all proteins identified by mass spectrometry, a principal component analysis (PCA) was performed on proteins having peptide spectrum matches in NDC80-GFP, but not control-GFP samples (schizont and gametocytes), and further excluding proteins that were annotated to be part of the ribosome (see **Fig. S5**). NA values were transformed into zero, indicating the absence of any peptide detection for a particular protein. PCA was done on ln(x) transformed unique peptide values using the ‘prcomp’ function, which is part of the R-package stats (v3.6.2). In a similar approach Spearman rank correlation was calculated for all proteins identified with both NDC80 and SPC25, using the ‘corr’ function, which is part of the R-package ggpubr (v0.2.5). Graphs were visualised using the R-package ggplot2.

### Comparative genomics of the NDC80 complex in Apicomplexa

#### Database

To increase the sensitivity for detecting highly divergent members of the NDC80 complex (NDC80, NUF2, SPC25, SPC24) in apicomplexan parasites, we constructed a large sequence database of (new) genomes and (meta)transcriptomes (see **Supplementary Table 3** for sources). This database consisted of 43 Apicomplexa and 8 Apicomplexa-affiliated lineages, including newly sequenced gregarines and ‘apicomonada’ (Janouskovec et al., 2019; Janouskovec et al., 2015; Mathur et al., 2019), and the non-parasitic apicomplexan *Nephromyces* (Munoz-Gomez et al., 2019). A set of 13 eukaryotes representative of a wider range of the eukaryotic tree of life were added for which the NDC80 complex presence/absence pattern was studied previously (van Hooff et al., 2017) For transcriptomes for which no gene predictions were available, ORFs were predicted using TransDecoder (Long Orfs algorithm: https://github.com/TransDecoder/TransDecoder).

#### Ortholog detection

To uncover an initial set of sequences orthologous to NDC80 complex subunits in our database we made use of previously established eukaryote-wide Hidden Markov Models (HMM) of the Calponin Homology domains of NDC80 and NUF2, and the RWD domains for SPC24 and SPC25 (Tromer et al., 2019; van Hooff et al., 2017). Although potentially informative, coiled-coil regions were avoided as they tend to have sequence similarities with (other) non-homologous coiled-coil proteins. For details on the strategy for finding orthologs, see previously established protocols (van Hooff et al., 2017). Briefly, HMM-guided hits were realigned using mafft (option:eins-i) (Katoh and Standley, 2013), modelled as HMM, and searched iteratively against the database until no new orthologs could be detected. Conspicuous absences were further inspected by iterative searches using sequences of closely related lineages, including lower stringency matches (higher E-values, lower bitscores) that had a similar length and coiled-coil topology. HMMs were modelled (hmmbuild) and iteratively searched (hmmsearch or jackhmmer: E-value<0.05, bitscore>25) using the HMMER package (v3.1b) (Johnson et al., 2010). SPC24 orthologues in Coccidia and Hematozoa were detected by comparing our custom-made eukaryote-wide HMMs with HMM profiles of automatically defined orthologous groups (OrthoFinder (Emms and Kelly, 2019)), using the secondary structure aware HMM-vs-HMM search algorithm HHsearch (Steinegger et al., 2019). Specifically, HMM profiles of the orthologous groups containing PBANKA_1442300 (*P. berghei*) and TGME49_212800 (*T. gondii)* were merged and the resulting HMM was searched against a dataset containing HMMs of the orthologous groups defined by OrthoFinder, a previously established set containing scop70, pdb70 and PfamA version 31.0 [7], and custom-made HMM of kinetochore proteins (Tromer et al., 2019; van Hooff et al., 2017).

Alignments were visualised and modified using Jalview (Waterhouse et al., 2009). Figure 6 was made using Inkscape (https://inkscape.org/).

## Acknowledgements

We thank Prof. Snezhana Oliferenko, The Francis Crick Institute, for stimulating discussions and advice on kinetochore proteins.

## Author contribution

RT conceived and designed all experiments. RT, MZ, ED and DB performed the experiments; RRS performed liver stage imaging; RP and RM performed super-resolution imaging; DJPF performed electron microscopy; ET and RW performed the bioinformatics and phylogenetic analysis; SA and KLR performed the ChIP-seq analysis; MZ, RT and ARB performed pulldown experiments, RP, MZ, AAH, and RT analysed the data; MZ, RP DSG, ET, RW AAH and RT wrote the manuscript and all others contributed to it.

## References

Akiyoshi, B., and K. Gull. 2013. Evolutionary cell biology of chromosome segregation: insights from trypanosomes. Open Biol. 3:130023.

Alushin, G.M., V.H. Ramey, S. Pasqualato, D.A. Ball, N. Grigorieff, A. Musacchio, and E. Nogales. 2010. The Ndc80 kinetochore complex forms oligomeric arrays along microtubules. Nature. 467:805–810.

Arnot, D.E., and K. Gull. 1998. The Plasmodium cell-cycle: facts and questions. Ann Trop Med Parasitol. 92:361–365.

Arnot, D.E., E. Ronander, and D.C. Bengtsson. 2011. The progression of the intra-erythrocytic cell cycle of Plasmodium falciparum and the role of the centriolar plaques in asynchronous mitotic division during schizogony. Int J Parasitol. 41:71–80.

Billker, O., V. Lindo, M. Panico, A.E. Etienne, T. Paxton, A. Dell, M. Rogers, R.E. Sinden, and H.R. Morris. 1998. Identification of xanthurenic acid as the putative inducer of malaria development in the mosquito. Nature. 392:289–292.

Bunnik, E.M., A. Venkat, J. Shao, K.E. McGovern, G. Batugedara, D. Worth, J. Prudhomme, S.A. Lapp, C. Andolina, L.S. Ross, L. Lawres, D. Brady, P. Sinnis, F. Nosten, D.A. Fidock, E.H. Wilson, R. Tewari, M.R. Galinski, C. Ben Mamoun, F. Ay, and K.G. Le Roch. 2019. Comparative 3D genome organization in apicomplexan parasites. Proc Natl Acad Sci U S A. 116:3183–3192.

Cheeseman, I.M. 2014. The kinetochore. Cold Spring Harb Perspect Biol. 6:a015826.

Ciferri, C., J. De Luca, S. Monzani, K.J. Ferrari, D. Ristic, C. Wyman, H. Stark, J. Kilmartin, E.D. Salmon, and A. Musacchio. 2005. Architecture of the human ndc80-hec1 complex, a critical constituent of the outer kinetochore. J Biol Chem. 280:29088–29095.

D’Archivio, S., and B. Wickstead. 2017. Trypanosome outer kinetochore proteins suggest conservation of chromosome segregation machinery across eukaryotes. J Cell Biol. 216:379–391.

Doerig, C., D. Chakrabarti, B. Kappes, and K. Matthews. 2000. The cell cycle in protozoan parasites. Progress in cell cycle research. 4:163–183.

Dorin-Semblat, D., C. Demarta-Gatsi, R. Hamelin, F. Armand, T.G. Carvalho, M. Moniatte, and C. Doerig. 2015. Malaria Parasite-Infected Erythrocytes Secrete PfCK1, the Plasmodium Homologue of the Pleiotropic Protein Kinase Casein Kinase 1. PLoS One. 10:e0139591.

Drechsler, H., and A.D. McAinsh. 2012. Exotic mitotic mechanisms. Open Biol. 2:120140.

Emms, D.M., and S. Kelly. 2019. OrthoFinder: phylogenetic orthology inference for comparative genomics. Genome Biol. 20:238.

Farrell, M., and M.J. Gubbels. 2014. The Toxoplasma gondii kinetochore is required for centrosome association with the centrocone (spindle pole). Cell Microbiol. 16:78–94.

Fennell, B.J., Z.A. Al-shatr, and A. Bell. 2008. Isotype expression, post-translational modification and stage-dependent production of tubulins in erythrocytic Plasmodium falciparum. Int J Parasitol. 38:527–539.

Ferguson, D.J., A.E. Balaban, E.M. Patzewitz, R.J. Wall, C.S. Hopp, B. Poulin, A. Mohmmed, P. Malhotra, A. Coppi, P. Sinnis, and R. Tewari. 2014. The repeat region of the circumsporozoite protein is critical for sporozoite formation and maturation in Plasmodium. PLoS One. 9:e113923.

Ferguson, D.J., F.L. Henriquez, M.J. Kirisits, S.P. Muench, S.T. Prigge, D.W. Rice, C.W. Roberts, and R.L. McLeod. 2005. Maternal inheritance and stage-specific variation of the apicoplast in Toxoplasma gondii during development in the intermediate and definitive host. Eukaryot Cell. 4:814–826.

Ferguson, D.J.P., N. Sahoo, R.A. Pinches, J.M. Bumstead, F.M. Tomley, and M.-J. Gubbels. 2008. MORN1 has a conserved role in asexual and sexual development across the apicomplexa. Eukaryotic cell. 7:698–711.

Francia, M.E., J.F. Dubremetz, and N.S. Morrissette. 2015. Basal body structure and composition in the apicomplexans Toxoplasma and Plasmodium. Cilia. 5:3.

Francia, M.E., and B. Striepen. 2014. Cell division in apicomplexan parasites. Nat Rev Microbiol. 12:125–136.

Gerald, N., B. Mahajan, and S. Kumar. 2011. Mitosis in the human malaria parasite Plasmodium falciparum. Eukaryot Cell. 10:474–482.

Guttery, D.S., M. Roques, A.A. Holder, and R. Tewari. 2015. Commit and Transmit: Molecular Players in Plasmodium Sexual Development and Zygote Differentiation. Trends in parasitology. 31:676–685.

Hoeijmakers, W.A., C. Flueck, K.J. Francoijs, A.H. Smits, J. Wetzel, J.C. Volz, A.F. Cowman, T. Voss, H.G. Stunnenberg, and R. Bartfai. 2012. Plasmodium falciparum centromeres display a unique epigenetic makeup and cluster prior to and during schizogony. Cell Microbiol. 14:1391–1401.

Iwanaga, S., S.M. Khan, I. Kaneko, Z. Christodoulou, C. Newbold, M. Yuda, C.J. Janse, and A.P. Waters. 2010. Functional identification of the Plasmodium centromere and generation of a Plasmodium artificial chromosome. Cell Host Microbe. 7:245–255.

Janouskovec, J., G.G. Paskerova, T.S. Miroliubova, K.V. Mikhailov, T. Birley, V.V. Aleoshin, and T.G. Simdyanov. 2019. Apicomplexan-like parasites are polyphyletic and widely but selectively dependent on cryptic plastid organelles. Elife. 8.

Janouskovec, J., D.V. Tikhonenkov, F. Burki, A.T. Howe, M. Kolisko, A.P. Mylnikov, and P.J. Keeling. 2015. Factors mediating plastid dependency and the origins of parasitism in apicomplexans and their close relatives. Proc Natl Acad Sci U S A. 112:10200–10207.

Janse, C.J., B. Franke-Fayard, G.R. Mair, J. Ramesar, C. Thiel, S. Engelmann, K. Matuschewski, G.J. van Gemert, R.W. Sauerwein, and A.P. Waters. 2006. High efficiency transfection of Plasmodium berghei facilitates novel selection procedures. Mol Biochem Parasitol. 145:60–70.

Janse, C.J., T. Ponnudurai, A.H. Lensen, J.H. Meuwissen, J. Ramesar, M. Van der Ploeg, and J.P. Overdulve. 1988. DNA synthesis in gametocytes of Plasmodium falciparum. Parasitology. 96 (Pt 1):1–7.

Johnson, L.S., S.R. Eddy, and E. Portugaly. 2010. Hidden Markov model speed heuristic and iterative HMM search procedure. BMC Bioinformatics. 11:431.

Katoh, K., and D.M. Standley. 2013. MAFFT multiple sequence alignment software version 7: improvements in performance and usability. Mol Biol Evol. 30:772–780.

Mathur, V., M. Kolisko, E. Hehenberger, N.A.T. Irwin, B.S. Leander, A. Kristmundsson, M.A. Freeman, and P.J. Keeling. 2019. Multiple Independent Origins of Apicomplexan-Like Parasites. Curr Biol. 29:2936–2941 e2935.

McKinley, K.L., and I.M. Cheeseman. 2016. The molecular basis for centromere identity and function. Nat Rev Mol Cell Biol. 17:16–29.

Munoz-Gomez, S.A., K. Durnin, L. Eme, C. Paight, C.E. Lane, M.B. Saffo, and C.H. Slamovits. 2019. Nephromyces Represents a Diverse and Novel Lineage of the Apicomplexa That Has Retained Apicoplasts. Genome Biol Evol. 11:2727–2740.

Musacchio, A., and A. Desai. 2017. A Molecular View of Kinetochore Assembly and Function. Biology (Basel). 6.

Pandey, R., S. Abel, M. Boucher, R.J. Wall, M. Zeeshan, E. Rea, A. Freville, X.M. Lu, D. Brady, E. Daniel, R.R. Stanway, S. Wheatley, G. Batugedara, T. Hollin, A.R. Bottrill, D. Gupta, A.A. Holder, K.G. Le Roch, and R. Tewari. 2019. Plasmodium condensin core subunits (SMC2/SMC4) mediate atypical mitosis and are essential for parasite proliferation and transmission. bioRxiv.

Petrovic, A., J. Keller, Y. Liu, K. Overlack, J. John, Y.N. Dimitrova, S. Jenni, S. van Gerwen, P. Stege, S. Wohlgemuth, P. Rombaut, F. Herzog, S.C. Harrison, I.R. Vetter, and A. Musacchio. 2016. Structure of the MIS12 Complex and Molecular Basis of Its Interaction with CENP-C at Human Kinetochores. Cell. 167:1028–1040 e1015.

Plowman, R., N. Singh, E.C. Tromer, A. Payan, E. Duro, C. Spanos, J. Rappsilber, B. Snel, G. Kops, K.D. Corbett, and A.L. Marston. 2019. The molecular basis of monopolin recruitment to the kinetochore. Chromosoma. 128:331–354.

Richmond, D., R. Rizkallah, F. Liang, M.M. Hurt, and Y. Wang. 2013. Slk19 clusters kinetochores and facilitates chromosome bipolar attachment. Mol Biol Cell. 24:566–577.

Roques, M., R.R. Stanway, E.I. Rea, R. Markus, D. Brady, A.A. Holder, D.S. Guttery, and R. Tewari. 2019. Plasmodium centrin PbCEN-4 localizes to the putative MTOC and is dispensable for malaria parasite proliferation. Biol Open. 8.

Saini, E., M. Zeeshan, D. Brady, R. Pandey, G. Kaiser, L. Koreny, P. Kumar, V. Thakur, S. Tatiya, N.J. Katris, R.S. Limenitakis, I. Kaur, J.L. Green, A.R. Bottrill, D.S. Guttery, R.F. Waller, V. Heussler, A.A. Holder, A. Mohmmed, P. Malhotra, and R. Tewari. 2017. Photosensitized INA-Labelled protein 1 (PhIL1) is novel component of the inner membrane complex and is required for Plasmodium parasite development. Sci Rep. 7:15577.

Schrevel, J., G. Asfaux-Foucher, and J.M. Bafort. 1977. [Ultrastructural study of multiple mitoses during sporogony of Plasmodium b. berghei]. J Ultrastruct Res. 59:332–350.

Sinden, R.E. 1983. Sexual development of malarial parasites. Adv Parasitol. 22:153–216.

Sinden, R.E. 1991a. Asexual blood stages of malaria modulate gametocyte infectivity to the mosquito vector--possible implications for control strategies. Parasitology. 103 Pt 2:191–196.

Sinden, R.E. 1991b. Mitosis and meiosis in malarial parasites. Acta Leiden. 60:19–27.

Sinden, R.E., E.U. Canning, R.S. Bray, and M.E. Smalley. 1978. Gametocyte and gamete development in Plasmodium falciparum. Proc R Soc Lond B Biol Sci. 201:375–399.

Sinden, R.E., E.U. Canning, and B. Spain. 1976. Gametogenesis and fertilization in Plasmodium yoelii nigeriensis: a transmission electron microscope study. Proc R Soc Lond B Biol Sci. 193:55–76.

Sinden, R.E., A. Talman, S.R. Marques, M.N. Wass, and M.J. Sternberg. 2010. The flagellum in malarial parasites. Curr Opin Microbiol. 13:491–500.

Solovei, I., L. Schermelleh, K. During, A. Engelhardt, S. Stein, C. Cremer, and T. Cremer. 2004. Differences in centromere positioning of cycling and postmitotic human cell types. Chromosoma. 112:410–423.

Steinegger, M., M. Meier, M. Mirdita, H. Vohringer, S.J. Haunsberger, and J. Soding. 2019. HH-suite3 for fast remote homology detection and deep protein annotation. BMC Bioinformatics. 20:473.

Sundin, L.J., G.J. Guimaraes, and J.G. Deluca. 2011. The NDC80 complex proteins Nuf2 and Hec1 make distinct contributions to kinetochore-microtubule attachment in mitosis. Mol Biol Cell. 22:759–768.

Suvorova, E.S., M. Francia, B. Striepen, and M.W. White. 2015. A novel bipartite centrosome coordinates the apicomplexan cell cycle. PLoS Biol. 13:e1002093.

Swedlow, J.R., K. Hu, P.D. Andrews, D.S. Roos, and J.M. Murray. 2002. Measuring tubulin content in Toxoplasma gondii: a comparison of laser-scanning confocal and wide-field fluorescence microscopy. Proceedings of the National Academy of Sciences of the United States of America. 99:2014–2019.

Tewari, R., D. Dorin, R. Moon, C. Doerig, and O. Billker. 2005. An atypical mitogen-activated protein kinase controls cytokinesis and flagellar motility during male gamete formation in a malaria parasite. Mol Microbiol. 58:1253–1263.

Tewari, R., U. Straschil, A. Bateman, U. Bohme, I. Cherevach, P. Gong, A. Pain, and O. Billker. 2010. The systematic functional analysis of Plasmodium protein kinases identifies essential regulators of mosquito transmission. Cell Host Microbe. 8:377–387.

Tromer, E.C., J.J.E. van Hooff, G. Kops, and B. Snel. 2019. Mosaic origin of the eukaryotic kinetochore. Proc Natl Acad Sci U S A. 116:12873–12882.

Vader, G., and A. Musacchio. 2017. The greatest kinetochore show on earth. EMBO Rep. 18:1473–1475.

Vaishnava, S., D.P. Morrison, R.Y. Gaji, J.M. Murray, R. Entzeroth, D.K. Howe, and B. Striepen. 2005. Plastid segregation and cell division in the apicomplexan parasite Sarcocystis neurona. J Cell Sci. 118:3397–3407.

van Hooff, J.J., E. Tromer, L.M. van Wijk, B. Snel, and G.J. Kops. 2017. Evolutionary dynamics of the kinetochore network in eukaryotes as revealed by comparative genomics. EMBO Rep. 18:1559–1571.

Verma, G., and N. Surolia. 2013. Plasmodium falciparum CENH3 is able to functionally complement Cse4p and its, C-terminus is essential for centromere function. Mol Biochem Parasitol. 192:21–29.

Verma, G., and N. Surolia. 2014. The dimerization domain of PfCENP-C is required for its functions as a centromere protein in human malaria parasite Plasmodium falciparum. Malar J. 13:475.

Waterhouse, A.M., J.B. Procter, D.M. Martin, M. Clamp, and G.J. Barton. 2009. Jalview Version 2--a multiple sequence alignment editor and analysis workbench. Bioinformatics. 25:1189–1191.

Wei, R.R., P.K. Sorger, and S.C. Harrison. 2005. Molecular organization of the Ndc80 complex, an essential kinetochore component. Proc Natl Acad Sci U S A. 102:5363–5367.

Westermann, S., D.G. Drubin, and G. Barnes. 2007. Structures and functions of yeast kinetochore complexes. Annu Rev Biochem. 76:563–591.

WHO. 2018. World malaria report 2018. World Health Organization.

Zeeshan, M., D.J. Ferguson, S. Abel, A. Burrrell, E. Rea, D. Brady, E. Daniel, M. Delves, S. Vaughan, A.A. Holder, K.G. Le Roch, C.A. Moores, and R. Tewari. 2019. Kinesin-8B controls basal body function and flagellum formation and is key to malaria transmission. Life Sci Alliance. 2.

