## Supplementary figures and images for "Real-time dynamics of *Plasmodium* NDC80 reveals unusual modes of chromosome segregation during parasite proliferation" (from another version of this preprint: version 4, 2020-02-27)

### Supplementary Figure

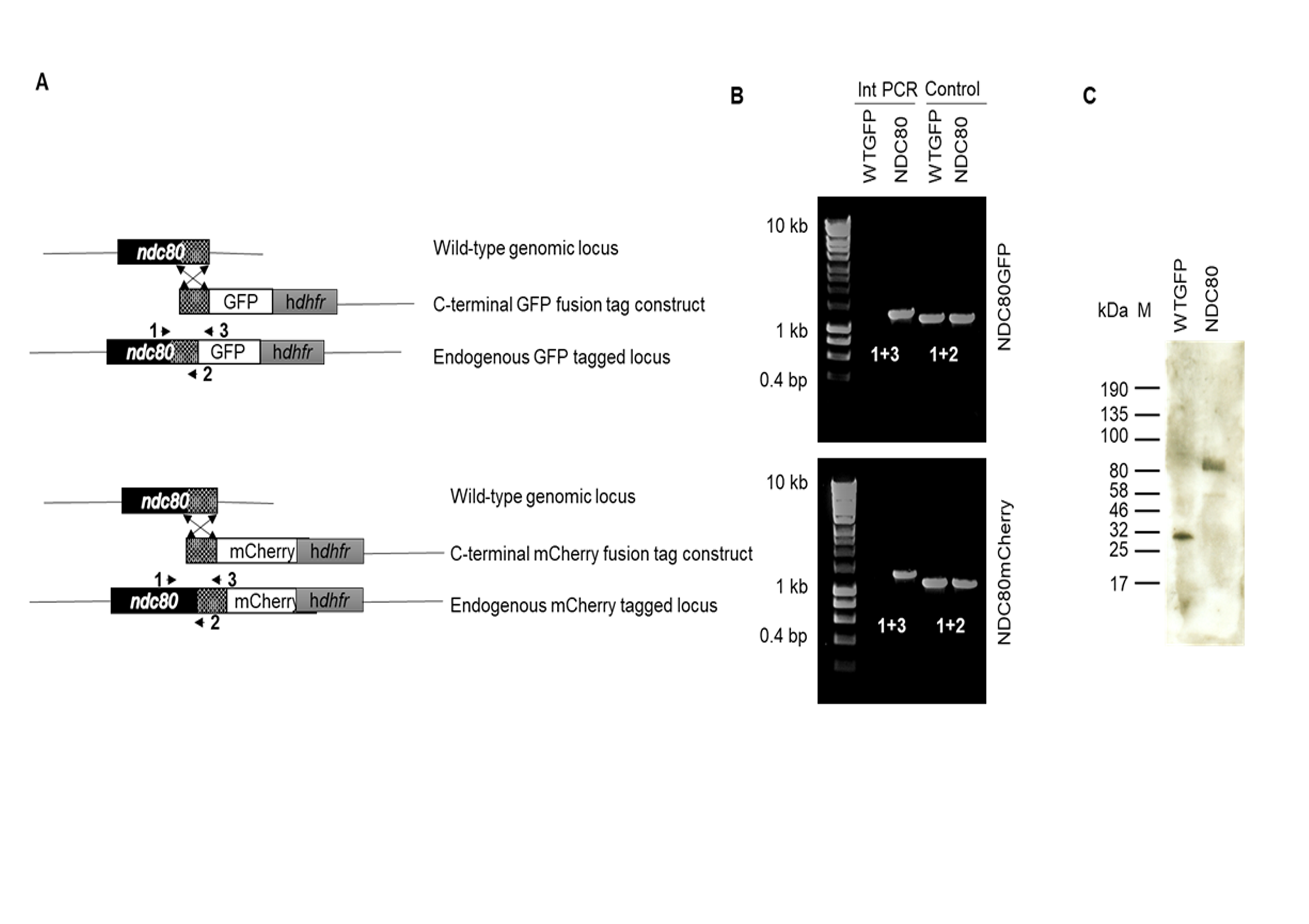

### Supplementary Figure

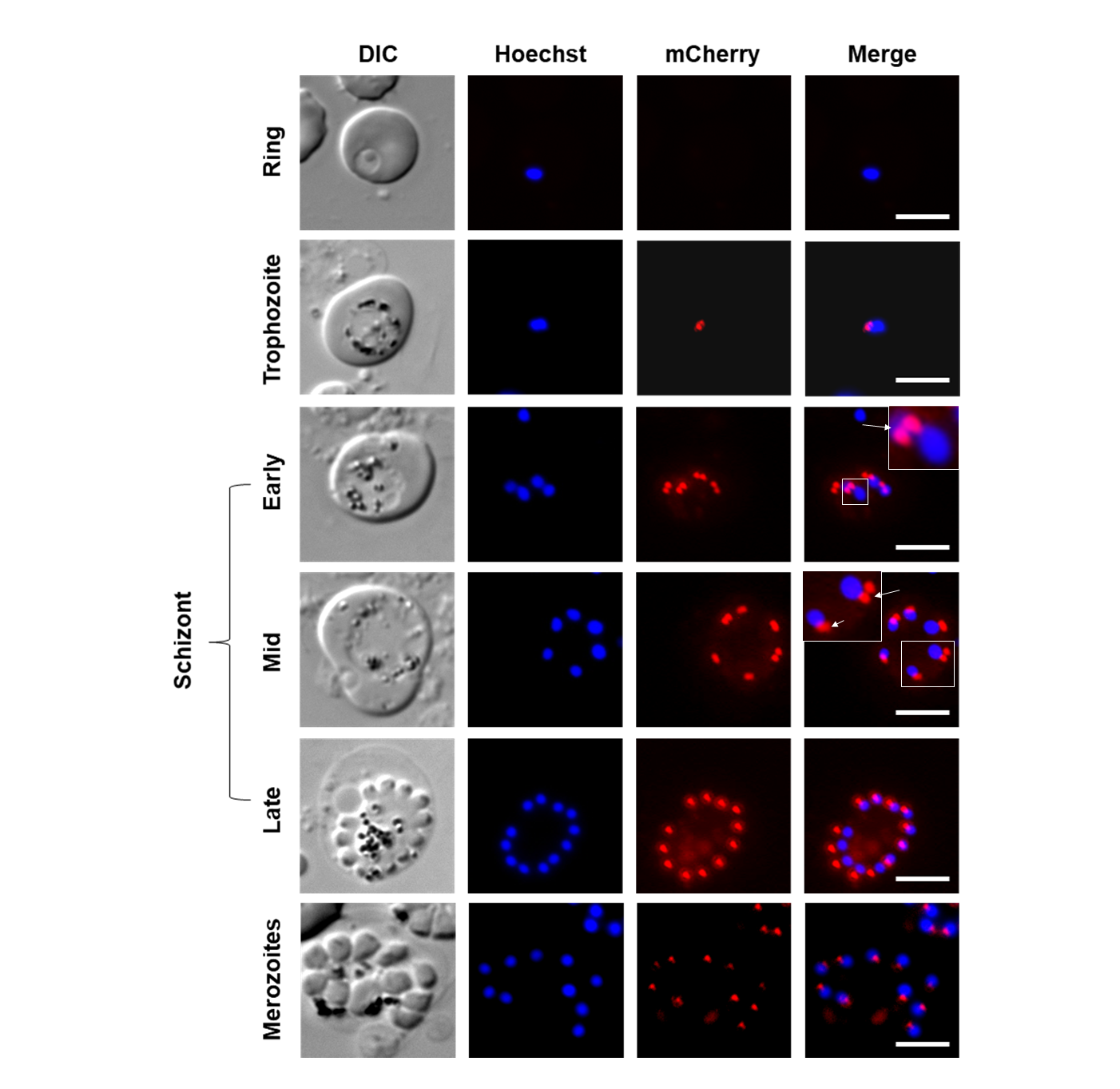

### Supplementary Figure

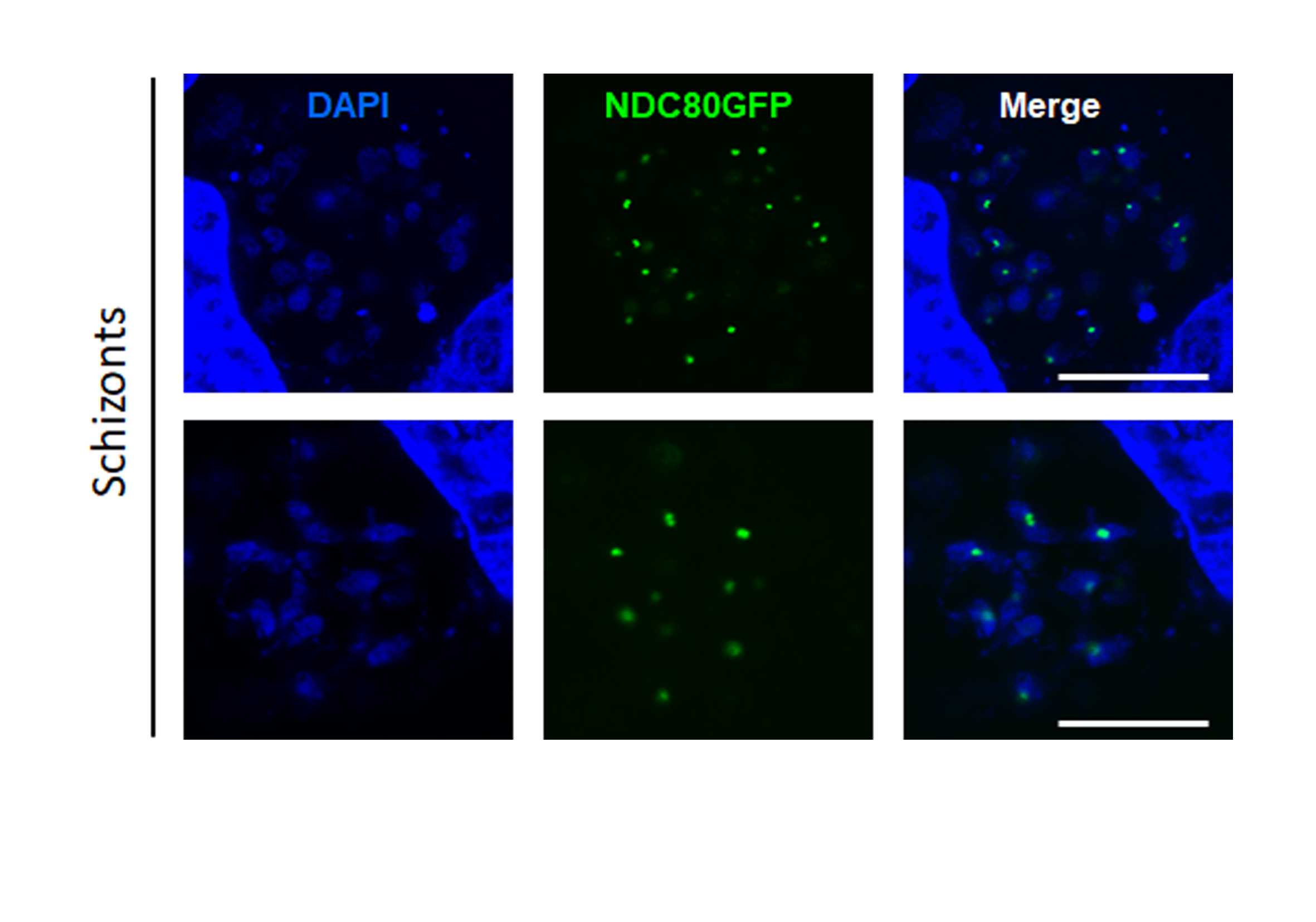

### Supplementary Figure

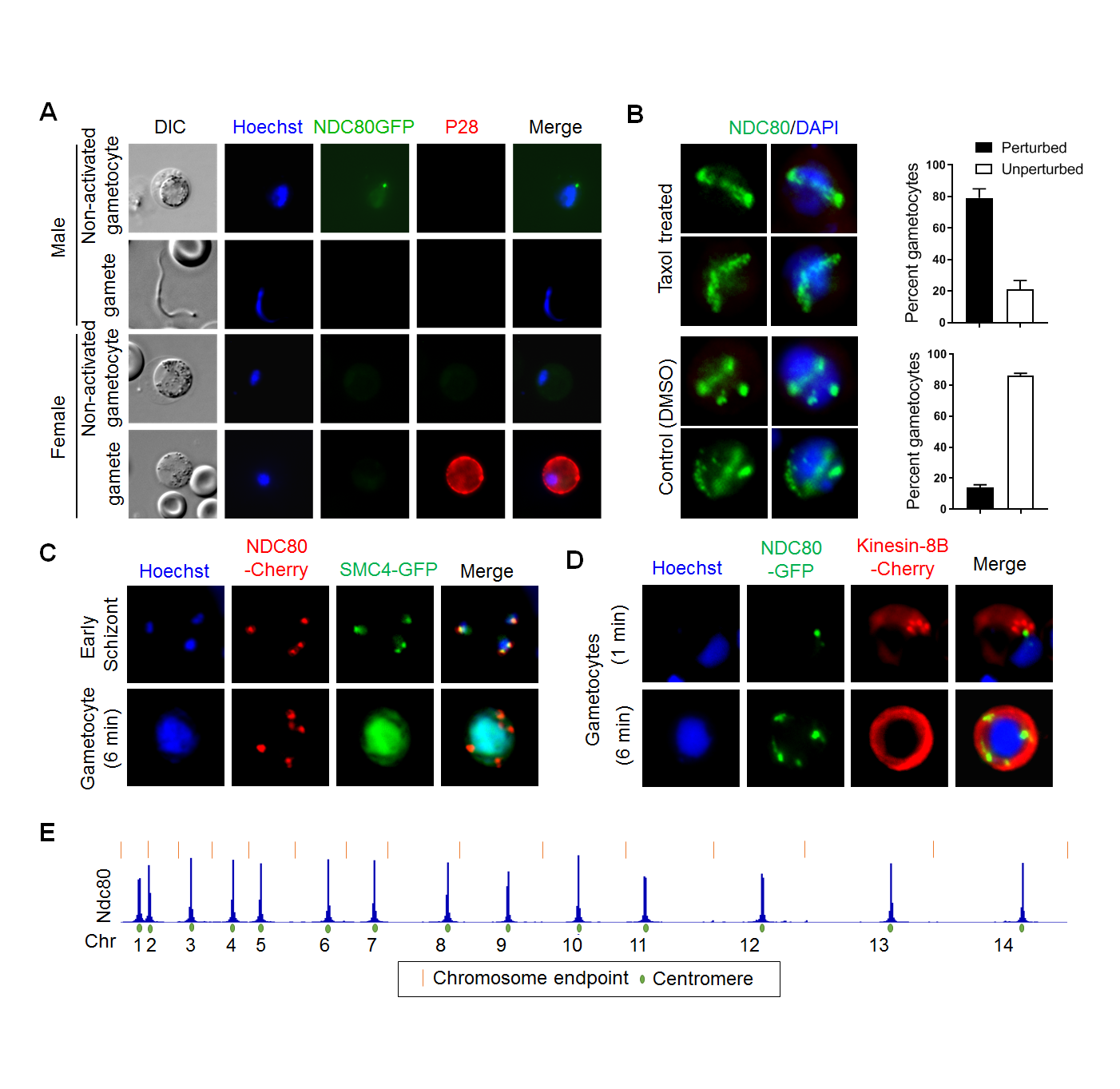

### Supplementary Figure

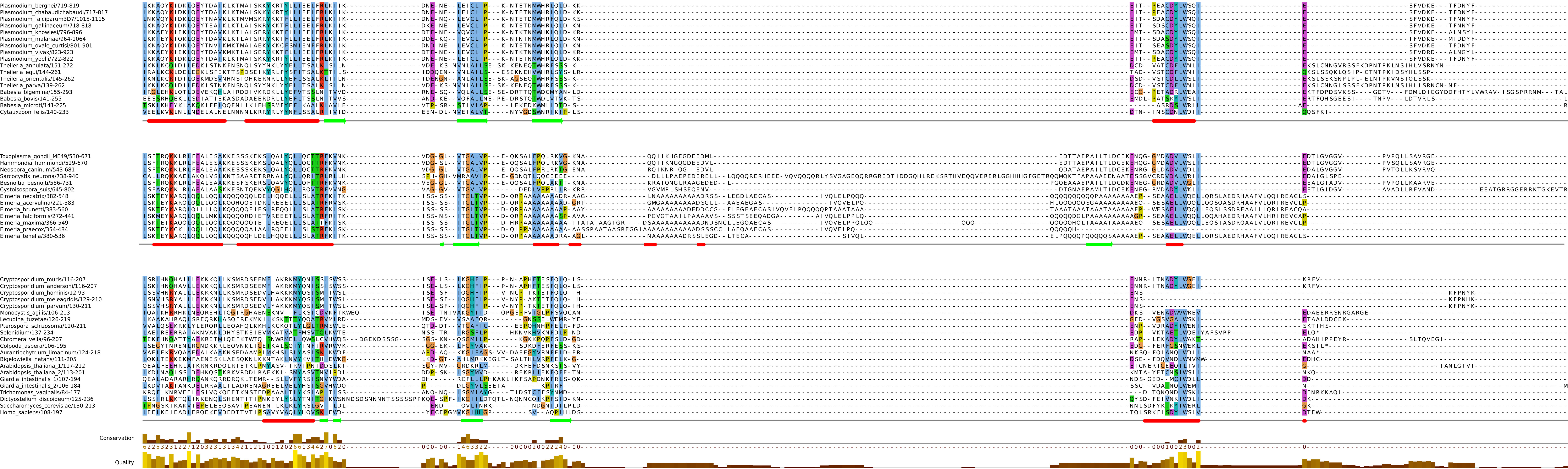
