## Supplementary Table for "Real-time dynamics of *Plasmodium* NDC80 reveals unusual modes of chromosome segregation during parasite proliferation" (from another version of this preprint: version 4, 2020-02-27)

| **Name** | **Sequence (5’ to 3’)** | **Notes** |
| --- | --- | --- |
| **T2591** | CCCCGGTACCGCGAATTCTAATAATAGGTTTAATC | KpnI site underlined |
| **T2592** | CCCCGGGCCCTTCAGTTACACTTTGATGTAAATTTTTATATAATTC | ApaI site underlined |
| **IntT259** | GATCTGGAAGAATCAATCAGAAAGAC |  |
| **ol492** | ACGCTGAACTTGTGGCCG | For GFP line |
| **mCherry** | TTCAGCTTGGCGGTCTGGGT | For mCherry line |

**Supplementary Table 1: Oligonucleotides used in this study**
